# Acquiring musculoskeletal skills with curriculum-based reinforcement learning

**DOI:** 10.1101/2024.01.24.577123

**Authors:** Alberto Silvio Chiappa, Pablo Tano, Nisheet Patel, Abigaïl Ingster, Alexandre Pouget, Alexander Mathis

## Abstract

Efficient musculoskeletal simulators and powerful learning algorithms provide computational tools to tackle the grand challenge of understanding biological motor control. Our winning solution for the inaugural NeurIPS MyoChallenge leverages an approach mirroring human skill learning. Using a novel curriculum learning approach, we trained a recurrent neural network to control a realistic model of the human hand with 39 muscles to rotate two Baoding balls in the palm of the hand. In agreement with data from human subjects, the policy uncovers a small number of kinematic synergies even though it is not explicitly biased towards low-dimensional solutions. However, by selectively inactivating parts of the control signal, we found that more dimensions contribute to the task performance than suggested by traditional synergy analysis. Overall, our work illustrates the emerging possibilities at the interface of musculoskeletal physics engines, reinforcement learning and neuroscience to advance our understanding of biological motor control.

## Introduction

Motor control, the art of coordinating muscles to produce intricate movements, is a marvel of biological intelligence. From the graceful dance of a ballerina to the dexterous manipulation of objects, these movements are a testament to the brain’s prowess of mastering numerous degrees of freedom ^1–5^ – which can take years of training and coaching to master, involving both explicit and implicit learning ^6,7^. Yet, understanding how the brain achieves skilled behavior remains one of the fundamental challenges in neuroscience. While significant strides have been made, much of the research has been confined to relatively simple behavioral tasks ^6–8^. Moreover, computational modeling of motor control and learning is usually limited to simplified models of the musculoskeletal systems ^9–13^. Of course, there are many good reasons to consider a simplified musculoskeletal system. While using these models has provided many fundamental contributions, a complementary approach is tackling motor skills with more realistic musculoskeletal models, which has so far remained out of reach ^5^.

Biomechanical simulators like OpenSim ^14^ have offered researchers many insights into musculoskeletal control ^5,15,16^. While deep reinforcement learning (RL) recently succeeded in training control policies for complex tasks ^17–23^, the computational cost for combining it with biomechanics simulators like OpenSim has been a bottleneck ^24,25^. MyoSuite ^26^, built on the efficient MuJoCo physics simulator ^27^, revolutionizes this space. It is not only vastly faster (up to 4000x) than its predecessors ^26^, but also introduces intricate object manipulation tasks like Baoding balls, which have long fascinated motor control researchers due to its demand on coordinated and dynamic fine motor skills ^3^.

In this article, we contribute to the ongoing dialogue between computational models and biological understanding in two significant ways. First, drawing inspiration from human skill learning, we introduce a novel learning approach: the Static to Dynamic Stabilization (SDS) curriculum. Our approach, SDS, won the first NeurIPS MyoChallenge for Baoding Balls ^28^; here we describe our solution in detail. Second, we analyze the learned policy, comparing and contrasting it with data from humans. We found that like biological agents ^1–5^, the SDS policy learns to operate in a reduced kinematic (pose) and dynamic (muscles) space, despite its generic architecture and the absence of pre-programmed simplifications. We found that the controller is robust to activity perturbations and that even low-variance dimensions still contain task-relevant signals, akin to Yan et al. ^29^. Furthermore, by considering additional object manipulation as well as control tasks, we found that the learned subspaces are task-dependent and do not generalize well to other tasks. Exploiting the biomechanical simulator, we also highlight that higher dimensional control spaces are needed to carry out the task than is suggested by traditional reconstruction analyses. Last but not least, we found lower tangling of the dynamics in the learned controller state space than in the action state space, akin to what Russo et al. ^30^ found for motor cortex vs. EMG dynamics. We discuss these results in light of the muscle synergy and skill-learning literature.

To our knowledge, SDS is the first example of an artificial agent successfully controlling a realistic musculoskeletal model of the hand in a skillful object manipulation task. Overall, our work showcases how simulation-based approaches can provide key insights into biological motor control, complementing experiments by providing a window into the representations underlying control, thereby allowing for studies across multiple scales of abstraction. With these advancements, we are better equipped to tackle deeper challenges in the realm of motor control, particularly the alignment between artificial and biological systems ^31^.

## Results

### Solution to the MyoChallenge

The Baoding balls task, as featured in MyoSuite, offers a rigorous testbed that captures the essence of contact-rich motor control challenges ^26,28^. It presents a biologically-realistic model of the human forearm, complete with a skeletal structure encompassing 23 joints that are actuated by 39 independent muscles, and two freely moving balls subject to physics (Figure 1A). The objective of the task is to maneuver the pair of balls in the hand, making them rotate in tandem along a circular trajectory. Maximum reward is achieved when the controller can guide the balls to follow a pair of moving targets (small spheres in Figure 1B).

**Figure 1.**
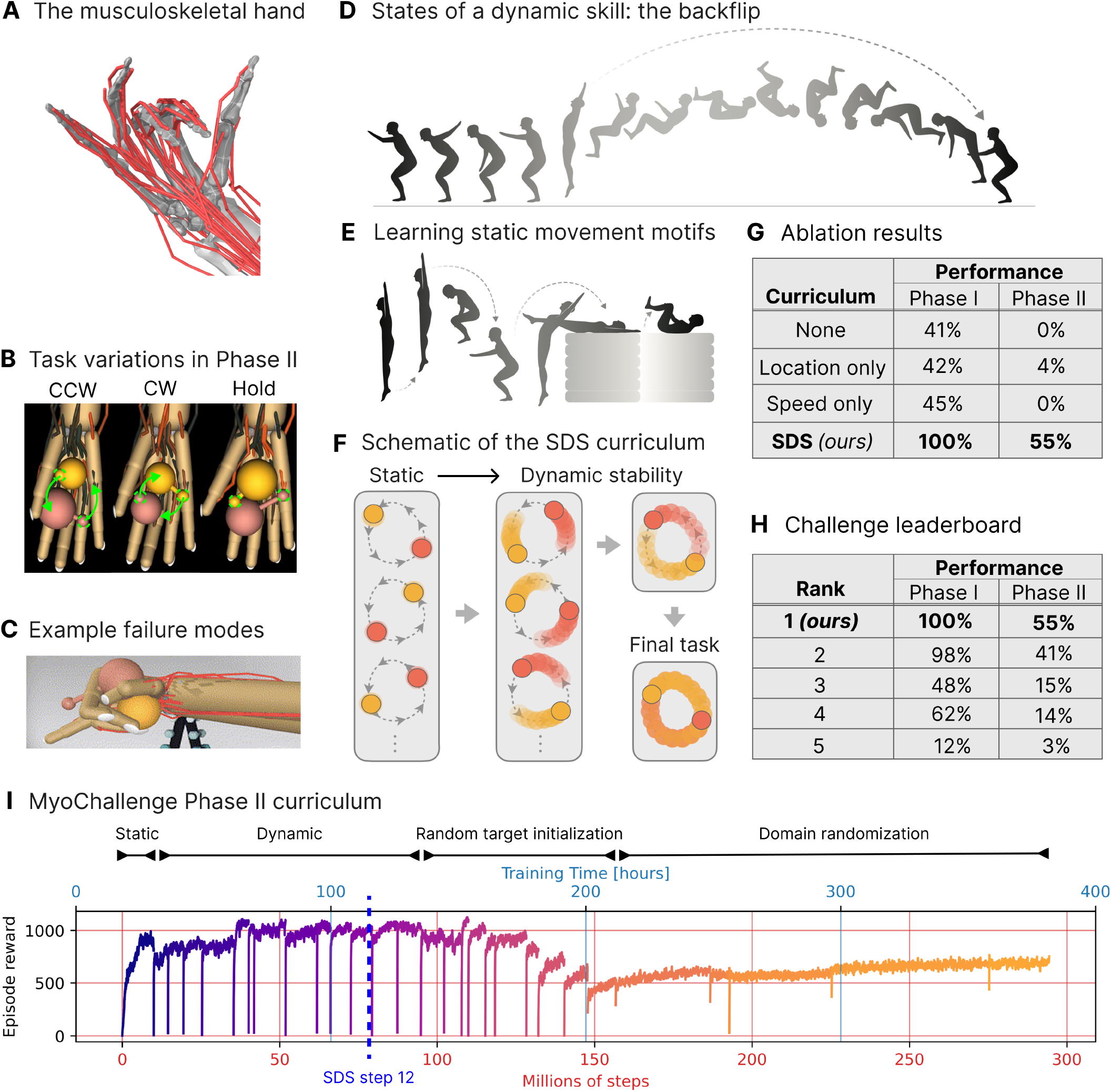
Definition of the Static to Dynamic Stabilization (SDS) curriculum, performance benchmarks and ablation study. **A** Visualization of the musculoskeletal hand in Mujoco ^26,27^. **B** Task variations include direction of rotation (counter-clockwise (CCW, left), clockwise (CW, middle), and hold (Hold, right)) along with domain randomization, i.e., varying initial locations of targets (dotted green circles) as well as varying ball size, mass, and friction. **C** Visualization of a problematic local optimum during curriculum learning. **D** Illustration of the states traversed during a backflip, a complex, whole-body skill performed by humans. **E** Steps involved in a recommended training routine. Illustrations in D and E by Julia Kuhl inspired by backflip training materials. **F** Schematic of our Static to Dynamic Stabilization (SDS) curriculum proposed to tackle the Baoding balls task by analogy to human skill learning (D-E). **G** Performance results from an ablation study demonstrate the necessity of the curriculum, especially in noisy environments with multiple conditions (Phase II of the MyoChallenge). Here and in H, performance is measured as the fraction of time steps in which the balls overlap with the targets. **H** Performance benchmarks and the MyoChallenge leaderboard (each row is a team). **I** Learning curve illustrating the 32 curriculum steps used to train the policy that achieved the top performance in Phase II of the MyoChallenge. The graph displays episode reward against training time and number of environment interactions. The reward is a dense signal which we designed to encourage the agent to follow the target trajectory of the Baoding balls without letting them fall. The initial *Static* and *Dynamic* periods correspond to the respective stages of the SDS curriculum. The latter part of the training focused on dealing with the misalignment between the initial position of the Baoding balls and the targets (*Random target initialization*) and with the variable environment physics (*Domain randomization*). Note that the maximum (possible) episode reward decreases during the *Random target initialization* period, as the targets do not overlap with the balls at the beginning of the episode. While the episode reward generally increases within a curriculum step, it exhibits a decreasing trend as more challenging settings are introduced during training. Our goal was to maximize performance on the (to-us) *u*nknown statistics of the Phase II testing. The dashed blue line corresponds to the twelfth step of the SDS curriculum, after which random target initialization and domain randomization are introduced; we later also use this network state for comparisons. See also Figure S1 and S2 and Table S1.

The MyoChallenge unfolded in two distinct phases. In Phase I, only *counter-clockwise* (CCW) rotations had to be achieved. In contrast, Phase II introduced several layers of complexity. Not only did it add the *clockwise* (CW) rotation and *stay still* (Hold) conditions, but it also necessitated complex decision-making right at the episode’s onset. The task’s condition was not inherently evident from the observed variables; instead, it had to be inferred by the agent. Adding to the challenge, the initial target position of the balls might not align with the actual position of the balls at the onset of an episode. This misalignment demanded strategic decisions to reach a high reward: whether to initiate the balls’ rotation in the reverse or forward direction, when to switch directions, decelerate, or maintain position. Further amplifying the task’s difficulty, Phase II introduced variability by randomizing parameters like the required targets’ rotation radius and speed, as well as the balls’ size, weight, and friction (Table 1). Each episode (in RL nomenclature, but more aptly described as ‘trial’ in Neuroscience) began with these parameters being randomly sampled from a predetermined range (Figure 1B).

**Table 1.**
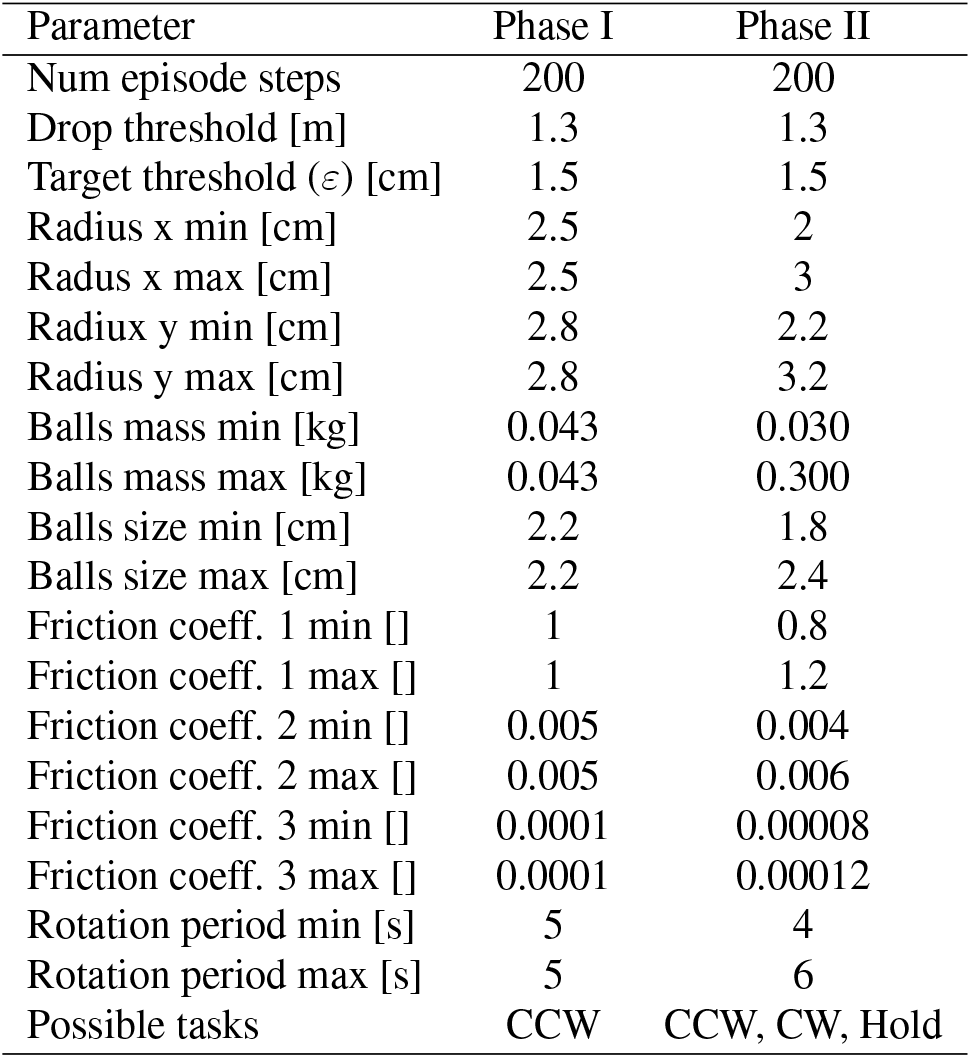
Parameters of the Boading Balls environments of Phase I and II.

| Parameter | Phase I | Phase II |
| --- | --- | --- |
| Num episode steps | 200 | 200 |
| Drop threshold [m] | 1.3 | 1.3 |
| Target threshold ($\varepsilon$) [cm] | 1.5 | 1.5 |
| Radius x min [cm] | 2.5 | 2 |
| Radius x max [cm] | 2.5 | 3 |
| Radius y min [cm] | 2.8 | 2.2 |
| Radius y max [cm] | 2.8 | 3.2 |
| Balls mass min [kg] | 0.043 | 0.030 |
| Balls mass max [kg] | 0.043 | 0.300 |
| Balls size min [cm] | 2.2 | 1.8 |
| Balls size max [cm] | 2.2 | 2.4 |
| Friction coeff. 1 min [] | 1 | 0.8 |
| Friction coeff. 1 max [] | 1 | 1.2 |
| Friction coeff. 2 min [] | 0.005 | 0.004 |
| Friction coeff. 2 max [] | 0.005 | 0.006 |
| Friction coeff. 3 min [] | 0.0001 | 0.00008 |
| Friction coeff. 3 max [] | 0.0001 | 0.00012 |
| Rotation period min [s] | 5 | 4 |
| Rotation period max [s] | 5 | 6 |
| Possible tasks | CCW | CCW, CW, Hold |

The evaluation criterion of the task measures the fraction of time steps within an episode when the balls successfully trace the desired trajectory marked by the moving targets (see Methods).<

### Reinforcement and curriculum learning

Recently, deep RL algorithms made strides in learning motor skills from scratch without a model of the environment dynamics ^18,19,23^. Such model-free RL algorithms have proven to scale robustly across increasingly complex environments. This robust scalability can be attributed to their effective use of powerful neural networks for processing high-dimensional data, alongside their high computational efficiency, broad applicability across tasks, and ease of ‘out-of-the-box’ implementation—advantages not as prevalent in more structured, model-based strategies. However, state-of-the-art model-free methods for RL faced significant challenges, which are evident when implementing them in the Baoding balls task: proximal policy optimization (PPO), a powerful model-free RL algorithm ^18^, combined with a recurrent neural network architecture (Figure S1) only reached 41% performance in Phase I and 0 % in Phase II (see Methods). The initial, sparse reward function proved insufficient for the agent to develop effective policies. Due to gravity and low friction, the balls easily rolled off the hand at the beginning of the episode, and the PPO agent was often not able to hold them at all. While adjusting the reward function to encourage long episodes provided an incentive to keep the balls on the palm (see Methods), the trained policy was highly susceptible to failure modes, like grabbing the balls to avoid them rolling off the hand (Figure 1C), preventing any meaningful behavior from emerging. These optimization challenges are inherent to model-free RL: as opposed to supervised learning, where the agent has access to the gradient of the task objective with respect to the control parameters, the RL agent must infer this gradient using the reward function. This makes learning less sample-efficient and more prone to exploration-exploitation failures. Hence, enhancing the model-free optimization process is crucial, which led us to focus on curriculum learning ^32–34^.

Drawing parallels to human skill learning can provide valuable inspiration ^35–38^. Consider the complex skill of performing a backflip (Figure 1D). For a novice, a backflip is a daunting sequence of (partially) unfamiliar states that must be seamlessly integrated into one fluid movement. Direct trial and error, or simply learning from one’s mistakes, can be a dangerous and inefficient approach. Athletes often undergo a structured training regimen: they first familiarize themselves with the specific bodily states required by the skill (Figure 1E). Once comfortable, they then piece these states together into a singular movement, initially slow and deliberate, eventually reaching full speed and fluidity.

Thus, inspired by coaching practice, we propose the Static to Dynamic Stabilization (SDS) curriculum for RL. Analogous to the athlete’s training, in SDS the RL agent is tasked to first learn to hold the balls statically in various configurations along the desired trajectory (Figure 1F, left). Once the controller can stabilize those states, the agent is gradually trained to dynamically transition between them, creating a continuous movement that mirrors the desired trajectory (Figure 1F, right). This curriculum not only addresses the exploration-exploitation dilemma but also provides the agent with a structured learning pathway, akin to the way humans approach some complex motor tasks. In the final stages of training, we also performed random target initialization and domain randomization, i.e. varying the task parameters (see Methods, Table 2). Overall, the SDS policy carried out 300 million environment interactions and it took almost 400 hours to train the model (Figure 1I). Due to the sequential nature of the curriculum, the training steps cannot be parallelized. The SDS policy achieved a 100% score in Phase I (Video 1), and 55% score in Phase II (Video 1), which is close to the maximum possible reward, since the initial positions of the balls and the targets do not overlap.

**Table 2.** Parameters of PPO and of the network architecture. The same parameters are used for both the actor and the critic. Some parameters changed starting from step 25 of the training curriculum.

| Parameter | Before step 25 | After step 25 |
| --- | --- | --- |
| Discount factor $\gamma$ | 0.99 | 0.99 |
| GAE $\lambda$ | 0.9 | 0.95 |
| Entropy regularization | 3.62109e-6 | 3e-5 |
| PPO clipping | 0.3 | 0.2 |
| Optimizer | adam | adam |
| Learning rate | 2.6e-5 | 2.5e-5 |
| Batch size | 2048 | 65536 |
| Minibatch size | 32 | 1024 |
| Max grad norm | 0.835671 | 0.8 |
| Num parallel envs | 16 | 16 |
| LSTM layer size | 256 | 256 |
| Layer 1 size | 256 | 256 |
| Layer 2 size | 256 | 256 |
| Activation | ReLU | ReLU |

We note that in the competition the sensory feedback was given in the form of joint angles and velocities, as it is more common in RL. We also tested if SDS could also learn from muscle lengths, velocities and forces, which is more akin to proprioceptive feedback ^39,40^. We found that a policy provided with such proprioceptive feedback and trained with SDS can achieve over 99% solved fraction in Phase I of the MyoChallenge (Figure S2). This control validates the SDS curriculum as a way to learn control policies also with more biologically realistic proprioceptive input.

### Performance benchmarks

To showcase the importance of the SDS curriculum, we performed an ablation analysis, systematically stripping away key components of the curriculum to observe the resultant impact on learning efficacy. We evaluated three alternative training procedures, maintaining the network architecture and RL algorithm, but devoid of the full SDS curriculum (see Methods). We evaluated the performance of PPO without a curriculum (*None*, Figure 1G); a curriculum that attempts to rotate the balls at full speed but departing from multiple initial locations (*Location only*, Figure 1G); and a curriculum that gradually increases the target speed but departing from a unique initial location (*Speed only*, Figure 1G). In the simpler Phase I, the performance drops by more than half upon removing either component (100% for SDS vs. 41-45% for *None, Location only, Speed only* curricula), whereas it all but entirely fails to learn anything in the more complex task of Phase II (55% for SDS vs. 0-4% for others). Thus the high performance of the SDS curriculum requires its two main components: learning multiple static configurations and gradually merging these configurations via increasingly faster dynamic trajectories.

To further contextualize our achievements, we juxtaposed our performance against those of the top contenders of the competition (Figure 1H). More than 40 teams took part and more than 340 entries were submitted ^28^. While most top solutions in the MyoChallenge incorporate RL with a curriculum, often complemented by reward shaping ^28^, it was our human-inspired SDS curriculum (Figure 1I, Static and Dynamic) that distinguished our approach and elevated our results above the competition. Furthermore, we reviewed the literature since the December 2022 challenge for new results: more recent exploration methods that do not use any kind of curriculum learning, such as generalized state-dependent exploration ^41^ and Lattice ^42^, marginally improve performance but fail to solve the task, successfully tracking less than 50% of the target ball trajectory (Table S1). This, further highlights the necessity of the curriculum learning approach.

To the best of our knowledge, our demonstration provided the first successful example of fully-learned musculoskeletal control in a skilled object manipulation task.

### Motor and muscle of synergies

Mastering the many degrees of freedom inherent in motor control, often referred to as *Bernstein’s problem* ^1–5^, is a central challenge in biological motor control. Having trained a policy network that solves the dexterous manipulation task with 39 muscles, we can interrogate the network state, muscle activations, and hand kinematics (Figure 2A). How does the artificial agent compare to the intricate behavior of human motor control?

**Figure 2.**
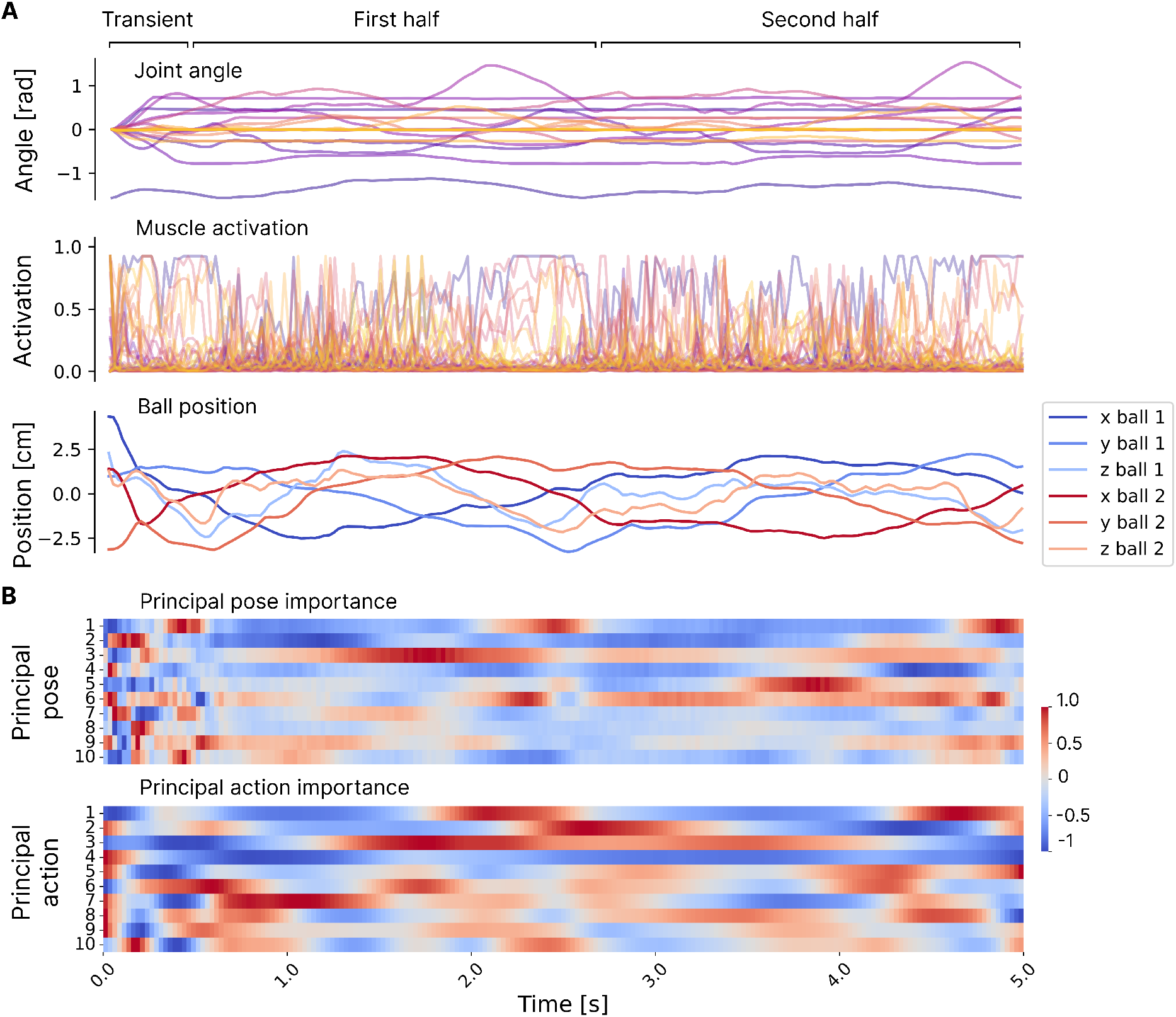
Kinematic and dynamic motifs within one full rotation of each Baoding ball for one episode of the SDS policy, after completion of the training curriculum. **A** Time series of the hand kinematics (top), the muscle activations (middle), and the position of the Baoding balls (bottom), during a full rotation of both balls (5 seconds). In the *transient* part of the trajectory, the hand catches the balls, which are initially slightly above the palm. In each of the subsequent halves of the trajectory, the position of the balls is swapped. **B** Relative importance of the 10 main principal components of the hand pose (top) and of the muscle activations (bottom), named *principal poses* and *principal actions*, as a function of the time step within a full rotation of both balls. The activations are averaged across all the episodes of the Baoding-SV dataset featuring a clockwise rotation (see Methods). Apart from the initial transient period, many components exhibit a periodic behavior, suggesting the emergence of coordination patterns which are repeated in the first and the second half of the rotation. Indeed, a full Baoding balls cycle is completed when the position of the two balls is swapped twice. See also Figure S3.

Classic motor control studies have illuminated a fascinating phenomenon: the vast realm of possible hand poses is not occupied by actual behavior, rather actual behavior is often distilled into a low-dimensional subspace ^1–5^. In other words, different joints appear coupled. Like others ^43,44^, we refer to the coordinated patterns of joint angles that span this low-dimensional subspace as *motor synergies*. Although researchers interpret motor synergies in various ways, it is widely accepted that dimensionality reduction techniques can be used to identify them ^3,5,43,45^. Here, we focus on analyzing these synergies and, in the discussion, interpret our findings in relation to the literature. Analogously, we refer to the basis set of muscle activation patterns as *muscle synergies*.

We hypothesize that similarly to biological agents ^1–5^, artificial agents also learn to operate in a reduced kinematic (pose) and dynamic (muscles) space. By projecting the hand poses (i.e., joint angles) and the policy’s actions (i.e., muscle activations) onto the principal axes, we can qualitatively gauge the significance of each principal component during the Baoding balls’ rotation cycle. Comparing the top principal poses and actions between the first and second half of the cycle, we see that the same principal component is reused to swap the positions of the Baoding balls (first row in both panels of Figure 2B), indicating that the SDS policy has effectively generalized across similar phases of the task.

We begin by comparing the SDS policy with human movement data from Todorov and Ghahramani ^3^, who used a CyberGlove to capture hand movements during object manipulation tasks, including the Baoding balls task. Their approach, using Principal Component Analysis (PCA) on the hand kinematics, sought to unveil the number of motor synergies at play. To get an upper limit they also considered a control task that instructed subjects to reach all joint limits. Echoing their methodology, we estimated the dimensionality of the movements generated by SDS (Figure 3A,B *Baoding*) and a policy for a control task called Hand Pose (Figure 3A,B *Control*), designed to span the maximum dimensionality of the hand in joint-space. Namely, to solve the hand-pose task, one needs to actuate the joints to random target hand postures (see Methods). The CyberGlove, however, does not record muscle dynamics, which is fully accessible in the MyoSuite simulator. Therefore, we can also analyze which activation patterns the SDS policy employs to control the hand, something that is more challenging to record in humans. For both tasks, we found that just a few synergies capture most of the variance in the posture (Figure 3A) and muscle space (Figure 3B). Note that if the posture space was being used uniformly, the cumulative variance plot would linearly increase. If it was spanned by a fixed combination of *N* independent primitives, then we should see a linear increase up to *N* principal components (PCs), where the cumulative variance becomes one. Given that this is not the case, we estimated the dimensionality with the same counting convention as in Todorov and Ghahramani ^3^, namely by averaging the number of dimensions needed to account for 85% and 95% of the variance (see Methods).

**Figure 3.**
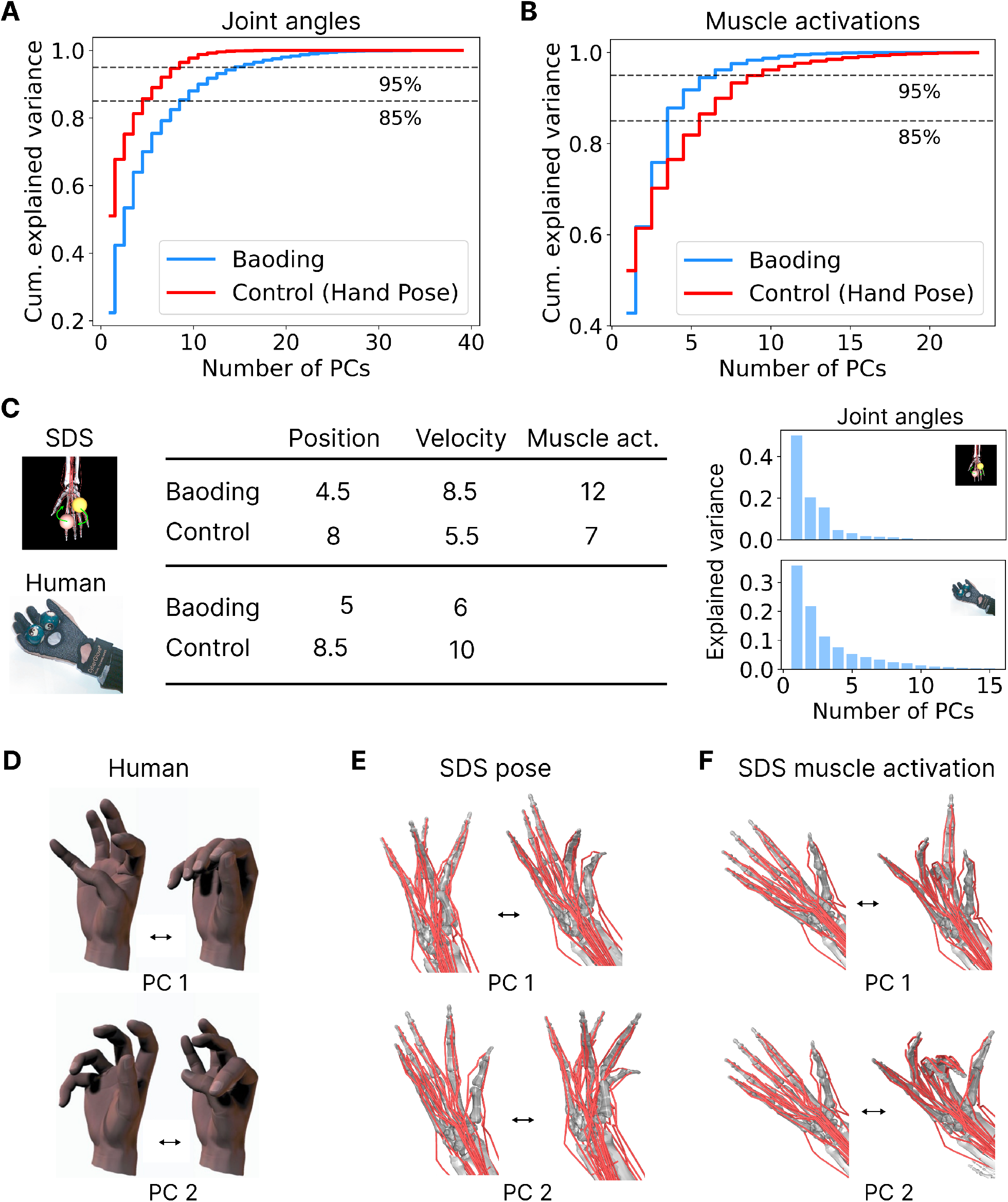
Dimensionality of the control policy and SDS policy in the Baoding balls task. **A** Cumulative explained variance of the PCs of the joint angles for an artificial agent trained to rotate two Baoding balls (SDS policy) and one trained to reach random poses (Control). For Baoding, the results in this plot (as well as in B and C) refer to the subset of episodes with counter-clockwise rotation direction. Also including the clockwise rotations gives similar results (Table 3). **B** Same as A, but the PCs are extracted from the agent’s actions, corresponding to the muscle activations. We also call those *muscle synergies*. **C** Left: Comparison between the number of independent degrees of freedom or *synergies* observed in the RL model (SDS) and experimental data (Human) ^3^ for the Baoding balls task as well as the control (hand/joint pose) task. The values are obtained by averaging the number of principal components necessary to explain 85% and 95% of the variance of the joint positions, the joint velocities and the muscle activations respectively (A, B). Right: Graphs of the explained variance for the first 15 principal components of the joint angles, for the SDS policy (top) and for a human subject (bottom, reproduced from Todorov and Ghahramani ^3^). **D** Pose variation corresponding to the first two principal components (PCs) of the hand poses recorded from humans rotating two Baoding balls (Figure adapted from Todorov and Ghahramani ^3^). **E** Same as D, but extracting the PCs from the hand poses of the SDS policy rotating the Baoding balls counter-clockwise. **F** Impact of applying the muscle activation patterns corresponding to the first two principal components of the control policy of the MyoChallenge competition for 15 steps (0.6 s), starting from an open hand position. See also Figure S4.

Our findings painted a compelling picture: the dimensionality of hand poses during Baoding balls rotation was lower than during the control task, both in our RL controller and in Todorov and Ghahramani ^3^’s experiments (Figure 3C, *Position* column). This result also validates the experimental design of the control task. Remarkably, despite the simulated agent being trained to maximize performance and not to match human behavior, the number of synergies in joint angle space computed for the simulated agent closely resembles the one measured for human subjects (Figure 3C, Position column), an observation also supported by the fraction of explained variance of individual components (Figure 3C, right). We also checked that our dimensionality estimates are robust to the dataset size (Figure S4).

**Table 3.** Detailed dimensionality estimation of the joint angular positions (Position), joint angular velocities (Velocity) and actions (Muscle activation). The values were extracted including the clockwise (CW), counter-clockwise (CCW) or both (Both) episodes of the Baoding-LV dataset (see Methods), and considering different thresholds of the explained variance.

| Exp. var. thresh. | Position |  |  | Velocity |  |  | Muscle activation |  |  |
| --- | --- | --- | --- | --- | --- | --- | --- | --- | --- |
|  | CW | CCW | Both | CW | CCW | Both | CW | CCW | Both |
| 0.85 | 4.0 | 3.0 | 4.0 | 6.0 | 6.0 | 6.0 | 9.0 | 9.0 | 9.0 |
| 0.95 | 6.0 | 6.0 | 7.0 | 10.0 | 11.0 | 11.0 | 15.0 | 15.0 | 15.0 |
| Average | 5.0 | 4.5 | 5.5 | 8.0 | 8.5 | 8.5 | 12.0 | 12.0 | 12.0 |

In contrast, a difference between artificial and human subjects emerged when analyzing the dimensionality of the angular velocities of the joints (Figure 3C, *Velocity* column). However, this may be expected. While the tasks on which we trained the artificial agents (Baoding and Control) resemble the experiments in terms of states to be visited (ball trajectory for Baoding and attainable hand poses for control), the instructions regarding the speed were different. The control policy attempts to reach the target pose as fast as possible, while for Baoding the rotation speeds are constrained by the task. In contrast, human subjects were instructed to solve the task at an (unspecified) comfortable speed ^3^, likely faster than the rotation speed of the SDS policy (4 - 6 s period).

Diving deeper, we performed an analysis that is harder with subjects: probing the dimensionality of the control signal in muscle-space. Strikingly, the dimensionality of the control policy in muscle-space showed a different pattern than in joint-space. Object manipulation demanded *more* degrees of freedom than pose reaching (12 vs. 7, Figure 3C). This finding introduces a critical, yet often overlooked perspective: assessing the complexity of control solely from motion capture observations might prove misleading ^5,46–49^. It is likely that this result, namely that the muscle-space dimensionality during free-hand movement is lower than during object manipulation, derives from the musculoskeletal structure of the hand, which has pairs of muscles antagonizing each other such that reaching desired hand positions requires low-dimensional muscle activation patterns. In contrast, robustly manipulating objects like two Baoding balls might require a more complex co-activation of antagonist muscles, leading to a higher effective muscle-space dimensionality. This interpretation is supported by the fact that the control policy outputs a sparser control signal than the Baoding policy (Figure 4C). Indeed, we can assess how many muscles a control policy recruits by computing, for every simulation time step, the number of muscles activated above a certain threshold. We found that the control policy, on average, only activates around 6 muscles out of 39 above a 5% activation threshold. This value is much larger for the Baoding policy, which activates, on average, over 17 muscles above the same 5% threshold. This indicates that the Baoding policy retains a residual activity for a large fraction of muscles, which is not present in the free reaching movements.

**Figure 4.**
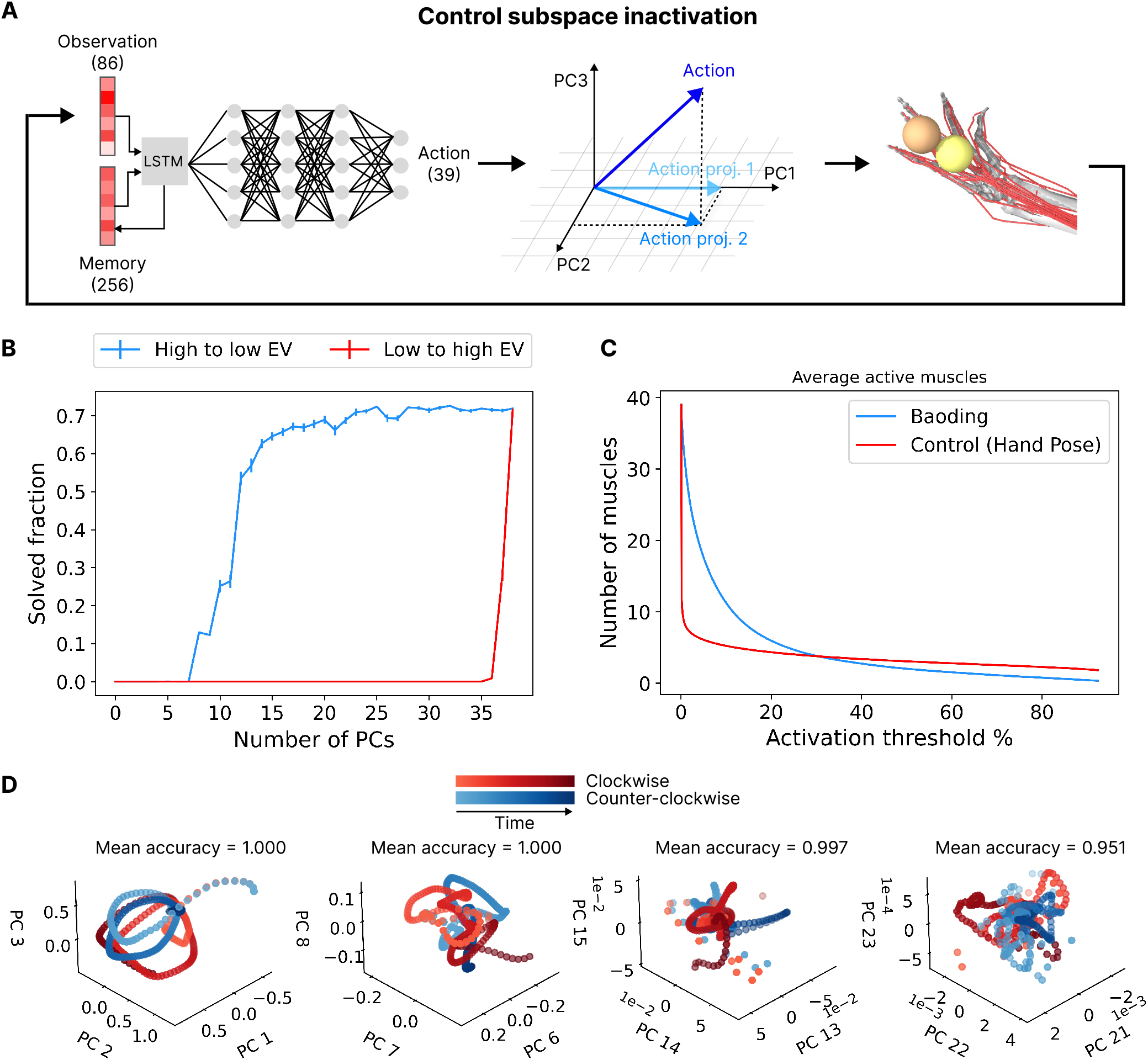
Task-relevance of the low-variance principal components of the SDS policy. **A** Illustration of the *control subspace inactivation procedure*. Before being transmitted to the environment, the action (i.e., pattern of muscle activations) is projected on the subspace spanned by a subset of the principal vectors and is therefore constrained to a lower dimensional space. The muscle activation pattern is applied to the hand model, which returns an observation to be processed by the policy network. **B** Average performance of the policy (± s. e. m. over 100 episodes) when the actions are projected onto progressively larger subspaces of the action space, defined by the principal components. Blue curve: the principal axes are added from the most to the least important in terms of explained variance. Red curve: the principal axes are added from the least to the most important. **C** Number of muscles active more than a threshold value, for variable threshold values. The control policy outputs a sparse control signal (the number of active muscles quickly drops for low activity threshold levels). For high activity thresholds, the control policy recruits slightly more muscles than Baoding. This indicates that the Baoding policy has a tendency to output intermediate activation values, while the control policy outputs extreme values (either fully active or not active at all). **D** Average trajectory of the hand pose (500 episodes per rotation direction, see Methods) when projected onto different 3-dimensional subspaces defined by progressively less important principal components. The mean accuracy score refers to the decoding via logistic regression of the task direction from each of the 1000 episodes considered to obtain the average trajectory (see Methods). While the accuracy decreases with the importance of the PCs, one can also reliably decode the task from the least important ones.

To better grasp the meaning of the PCs of the hand pose, we can visualize them as shifts from the mean pose. Comparing the first two PCs computed from a human subject (Fig 3D) and from the SDS policy (Figure 3E), the SDS policy’s movements appear more rigid and localized to a subset of fingers. Furthermore, we can visualize the effect of individual PCs of the muscle activations by repeatedly applying them as actions, starting from an open hand position (Figure 3F). In the SDS policy, the first principal action causes a flexion of the middle finger and the thumb, and a wrist movement, while the second principal action orchestrates a flexion pattern of the thumb away from the palm and the other fingers towards it. The SDS policy primarily engages four of the five fingers (see videos 1 and 1). In particular, the policies of Phase I and of Phase II seem to not involve the middle and the ring finger, respectively. This contrasts with Todorov and Ghahramani ^3^‘s findings, where the first two PCs of the hand poses influenced all fingers (Figure 3D). Why are those patterns different, when the dimensionality is comparable? Unlike biological agents, the artificial policy has been optimized exclusively for the Baoding task, which might explain the more localized and rigid movements. For the distribution of ball sizes, engaging four fingers might be sufficient, and gives rise to a similar motor synergy dimension. Discrepancies in the biomechanical model could also be at play.

Overall, we found that like biological agents ^1–5^, artificial agents learn to operate in a reduced kinematic (pose) and dynamic (muscles) space.

### Task relevance of the low-variance PCs

It is tempting to categorize the high- and low-variance PCs as the task-relevant and task-irrelevant muscle synergies, respectively. Indeed, projecting the muscle activations onto the first 15 (out of 39) PCs accounts for over 95% of the variance of the control signal (Figure 3B). Yet, the role of low-variance (muscle) PCs in mastering the skill is unclear.

Indeed, we argue that the importance of muscle synergies is more meaningfully evaluated based on their impact on the task performance, rather than their contribution to the reconstruction of the control signal. The task performance can of course be evaluated in the biomechanical simulator ^5,46–49^. While inactivating specific muscle synergies is currently impossible experimentally, it is a straightforward procedure in our computational model. With an experiment akin to selective modulation of ensembles of neurons via optogenetics, we removed the component of the control signal, which lies on a specific subspace of the muscle activation space (Figure 4A). By projecting the muscle activation pattern output by the policy onto the subspace spanned by a subset of the principal directions of variability, we forced the control signal to lie in a lower-dimensional space (namely, the lower-dimensional space that captures the largest amount of variability). In this way, we could measure how the task performance varied as a function of the dimensionality of the enforced space (see Methods). We call this procedure *control subspace inactivation* (CSI). Adding one principal component after the other in decreasing order of explained variance (blue curve in Figure 4B) revealed how many directions of variability contribute to successfully rotating the Baoding balls. We found that twelve PCs out of 39 retain 50% of the task performance, which only saturates after 25 components are allowed. Compared to the dimensionality estimation obtained by reconstructing the control signal (Figure 3B), this estimation in terms of task performance returns a larger and, arguably, more meaningful value. In particular, even the components that contribute less than 5% in explaining the variance of the muscle-activation space play a decisive role in solving the task. Conversely, as expected, the high-variance PCs are crucial for task performance (red curve in Figure 4B), as just removing a few of them causes the *solved fraction* to quickly drop to zero.

Comparing activity across different conditions of the task can additionally shed light on the low-variance PCs of hand kinematics (clockwise or counter-clockwise rotation in Figure 4D). We found that low-variance PCs of the hand pose are task-dependent, as they retain discriminative power regarding what task is being performed (clockwise or counter-clockwise rotation in Figure 4D). This result is consistent with the observations of Yan et al ^29^, who showed evidence for task-relevance even in low-variance PCs for hand kinematics. Highly task-specific kinematic synergies suggest that it might not be possible to create a common, low-dimensional control subspace that works across tasks. To further investigate this question, in the next subsection we performed CSI across tasks, to show that kinematic or muscle spaces are indeed highly-task dependent.

### Task dependence of the muscle synergies

We have found that a policy trained with RL finds a low-dimensional kinematic and muscle space. It is unclear whether this emerging dimensionality reduction indicates the existence of redundant control dimensions that would allow a policy to confine itself to a reduced space for any motor task, or whether the dimensionality reduction is instead task-specific. To disentangle these possibilities, we considered three additional motor control tasks: Hand Reach, Reorient and Pen, featured in MyoSuite (see Methods). In Hand Reach, a policy has to control the MyoHand to reach five target points with the hand’s fingertips. Unlike the control task, which requires targeting specific angles for each joint, this task involves guiding only the fingertips to random targets. For Reorient and Pen, a policy has to control the MyoHand and rotate an object (a die and a pen, respectively) to achieve a target orientation. We also considered the policy obtained at step 12 of the SDS curriculum (*SDS step 12*), which achieved 100% *solved fraction* in the Baoding balls rotation, both CW and CCW, before any variability (rotation speed and radius, ball mass, size, and friction) was introduced (dashed blue line in Figure 1I). First, we evaluated the similarity between pairs of tasks in the kinematic and muscle space. For each of the six policies, we collected a dataset of 1000 episodes and extracted the principal components of the hand kinematics and of the muscle activations, defining a task-specific subspace. We then projected the kinematics and the muscle activations of the SDS step 12 policy (Figure 5A,B) and of the final SDS policy (Figure 5D,E) onto these task-specific subspaces. The PCA of the hand kinematics (Figure 5A,D) confirms the intuition that the final SDS policy and the SDS step 12 policy are the most similar, while the Hand Pose and Hand Reach policies are the most dissimilar from the SDS step 12 policy and the final SDS policy. In the muscle-activation space, the SDS step 12 and the final SDS are more different than in the kinematics space, and the clear hierarchy among the other tasks also disappears (Figure 5B, E). Thus, to accurately reconstruct the muscle dynamics of SDS one needs more than 20 muscle synergies from other tasks. However, this does not answer whether one could actually achieve the task. To tackle this, we take advantage of the biomechanical simulator.

**Figure 5.**
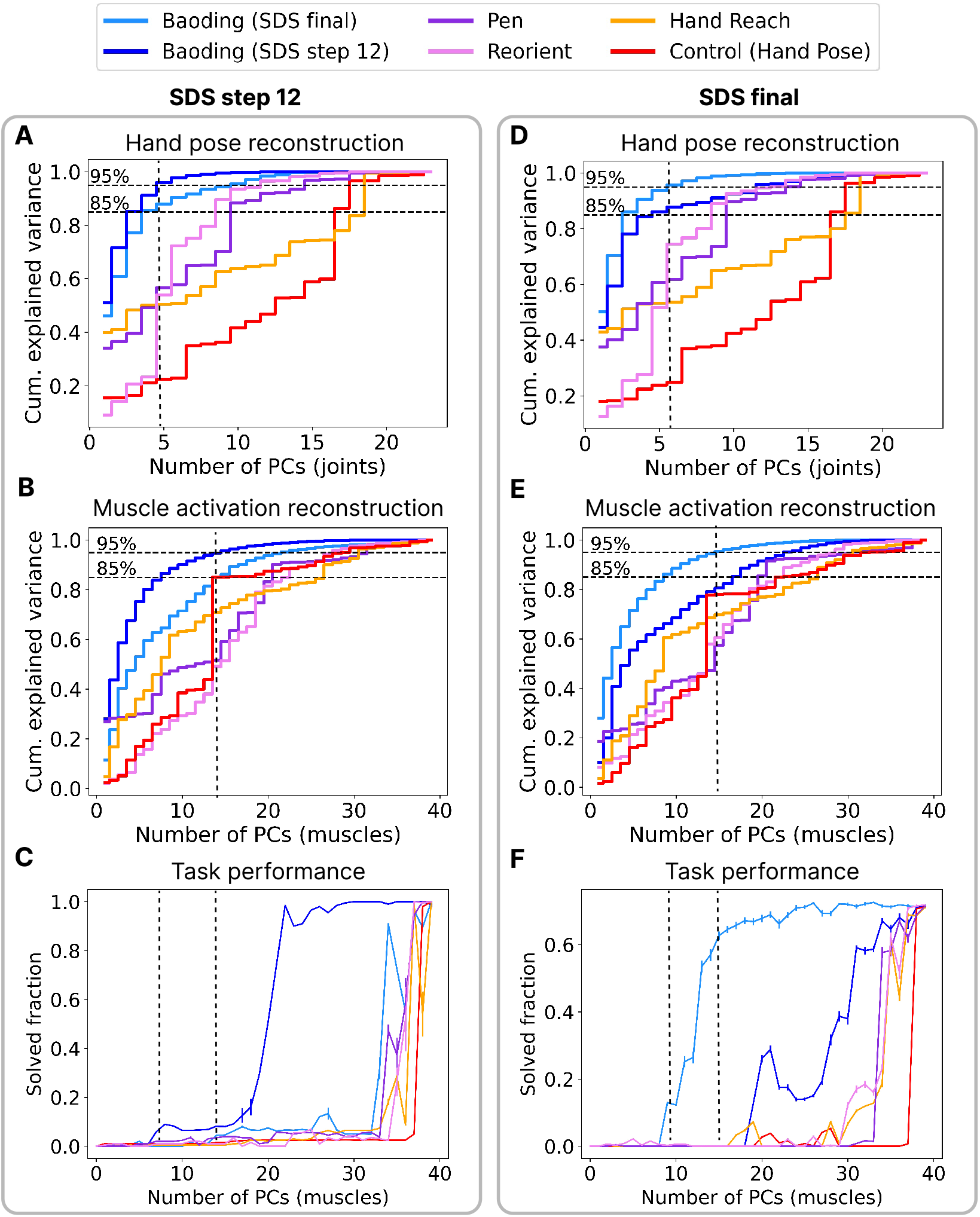
Transfer of muscle synergies from other tasks to Baoding. **A** Cumulative explained variance of the PCs of the joint angles for the SDS policy after 12 curriculum steps, performing counter-clockwise rotations. The explained variance is computed both on the principal components extracted from the same policy (SDS step 12) and from other tasks (final SDS, Hand Pose, Hand Reach, Pen, Reorient). The dashed vertical lines indicate the number of components extracted form the same task necessary to reach 95% explained variance. **B** Same as A, but the PCs are extracted from the agent’s actions, corresponding to the muscle activations. **C** *Control subspace inactivation* applied to the SDS step 12 policy, using PCs extracted from the policy itself and from those trained on the other tasks. The dashed lines indicate the dimensionality estimation based on 95% explained variance when reconstructing the hand pose and the muscle activations. **D-F** Same ad A-C, but for the final SDS policy. See also Figure S5.

Performance is a better metric for determining whether a common low-dimensional subspace can be reused to solve multiple tasks. This can be achieved by extracting the PCs of each task and using them as a basis for the inactivated sub-space when applying CSI on a different task. We performed this analysis on the SDS step 12 policy (Figure 5C) and on the final SDS policy (Figure 5F). We can see how, remarkably, 30 or more dimensions extracted from a different task are necessary to achieve high performance for both the SDS step 12 policy and the final SDS policy. Interestingly, while the kinematics of SDS step 12 and of final SDS are similar, the two tasks require surprisingly different muscle synergy spaces. A high number of muscle synergies is necessary to achieve good task performance also when extracting them from multiple tasks at the same time, although (as expected) they tend to transfer better to Baoding when all the tasks (Hand Pose, Hand Reach, Pen and Reorient) are considered at the same time (Figure S5). Taken together, these results indicate that, unless one uses more than 30 dimensions, the muscle synergies discovered by a policy trained on one task do not constitute a useful subspace to solve a different task. This estimate of at least 30 dimensions is close to the dimensionality of the original space and much bigger than previous estimates based on dimensionality reduction ^43,44^.

### Analysis of the SDS policy’s computation

The policy network, with its Long-Short Term Memory (LSTM) layer and two fully connected layers (Figure S1), offers a lens into how information is transformed from the hand’s proprioceptive state to the output control signal in terms of muscle activations. We embedded the activity of the policy network into a 3D space with UMAP (Figure 6A, see Methods). We also quantified the degree to which the state spaces of the two tasks are entangled using Russo et al.’s tangling metric *Q* (see Methods).

**Figure 6.**
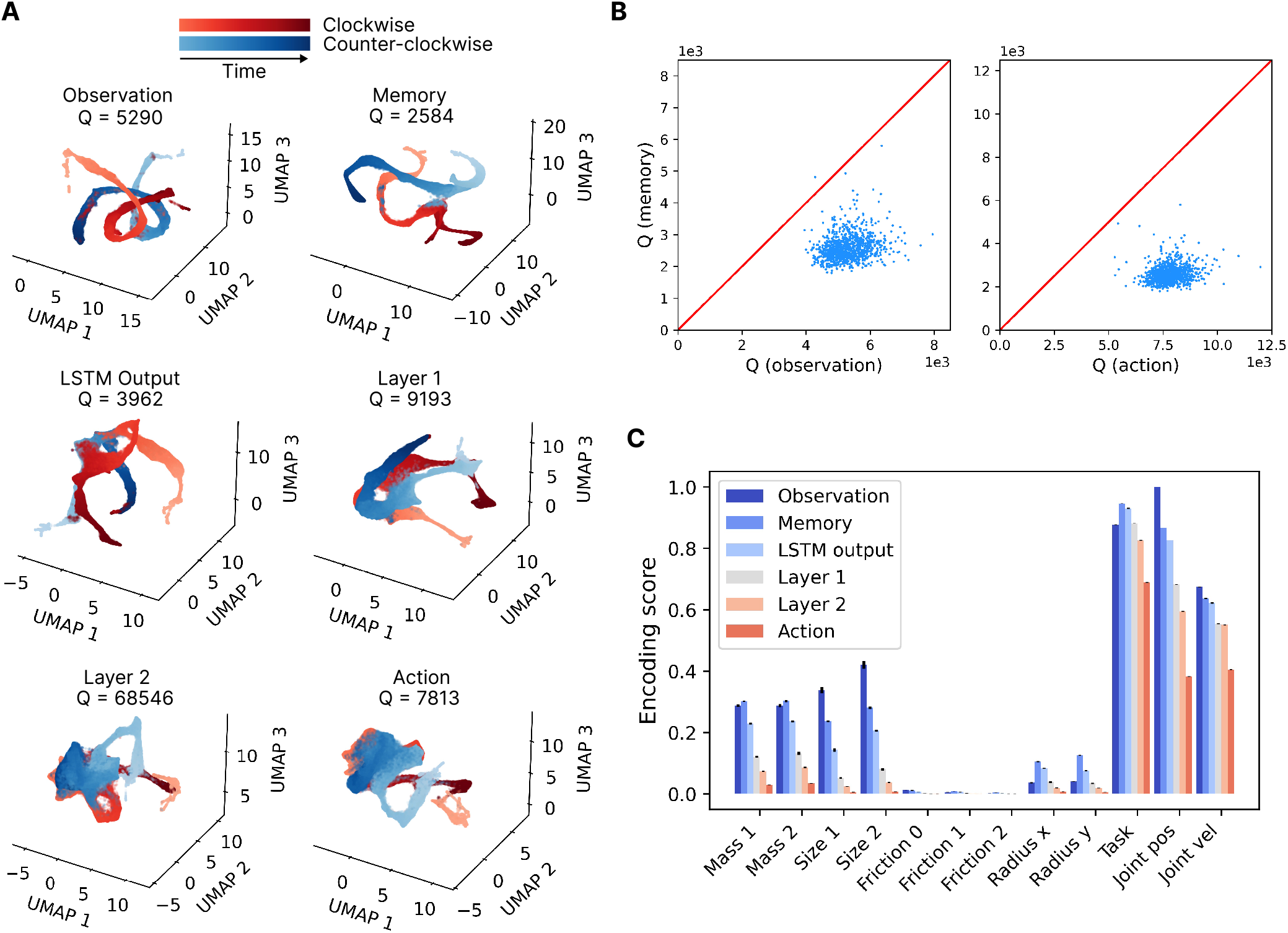
Population activity of the SDS policy network. **A** UMAP embedding of the population activity of all the layers of the policy network (Figure S1), extracted from the Baoding-SV dataset (see Methods). The rotational dynamics and the separation between clockwise and counter-clockwise rotation progressively disappear along the processing hierarchy. The average tangling metric *Q* is displayed (see Methods). **B** Pair-wise comparison of the tangling of the population activity in the memory layer versus the observation (left) and the action space (right). Each point indicates the average tangling of one episode (see Methods). The memory layer consistently untangles the input observation, while the deeper layers transform it into more tangled actions. **C** Decoding analysis of the policy network, showcasing where task-relevant quantities are best encoded in each layer. Each time point is separated by 25 ms. The error bars indicate the standard deviation across 5-fold cross-validation.

First, note that in the deeper layers (layer 1 and layer 2) and in the output (action), the activity of the two tasks (blue and red) is more tangled than in the input (observation) and in the early layers (memory and LSTM Output). Furthermore, the manifolds of the two tasks are more separated in the memory subspace than in the observation subspace (compare top-right with top-left), suggesting that the system distills the information from the observation space into two separate state spaces, effectively separating states that are similar in observation space but require different motor plans. Then, the state spaces merge back together towards the action space, as the system recruits similar motor synergies independently of the rotation direction (Figure 6A). Note that, unlike the tasks in the previous section, in which a different neural network was trained independently in each task, in this case the same neural network was trained to solve both the CW and CCW tasks, likely favoring the reusability of common synergies across tasks.

The population activity of the Memory layer of the LSTM block presents the least tangled trajectories in the policy network. In particular, it is consistently less tangled than the action and observation dynamics (Figure 6B) when carrying out the analysis in a per-episode manner. This result qualitatively resembles Russo et al.’s insight that motor cortical dynamics avoid tangling more than muscle dynamics (EMG) ^30^.

Finally, we sought to determine if and where in the network different task-relevant quantities are encoded (Figure 6C). While certain variables, like joint angular positions, can be directly decoded from the observation, others, that are not part of the observation space, like the physical properties of the balls, are better decoded from the LSTM layer’s memory layer. This suggests that the SDS policy accumulates data over time to form a representation of the system’s non-observable state, which then conditions the control signal; several variables cannot be well decoded suggesting that they are implicitly represented.

## Discussion

The intersection of efficient musculoskeletal simulators and powerful learning algorithms heralds a new era of understanding in the realm of biological motor control. By leveraging these models, we can delve deeper into the core principles of biological skill learning. This approach can not only offer insights into how animals, including humans, learn and execute complex motor tasks, but also generate both behavioral and neural predictions. The SDS curriculum, inspired by human learning, emphasizes the significance of reinforcement and curriculum learning in training motor control policies for complex tasks. This showcases the potential of RL in developing high-fidelity sensorimotor models and provides a platform to juxtapose artificial and biological control systems. Such comparisons, as we have demonstrated, can yield insights into emergent movement dynamics, offering a fresh perspective on motor learning and control.

Yet, as we push the boundaries of what is possible with digital simulations in deep RL, we are confronted with a deeper, more fundamental challenge: model alignment in sensori-motor systems. How do we bridge the gap between artificial systems and the intricate, naturalistic movements of a living organism? While supervised learning offers some solutions, allowing us to enforce specific movement repertoires, as demonstrated in previous models of the motor ^50^, visuo-motor ^51^ and proprioceptive system ^40,52^, RL presents a unique conundrum. Without explicit guidance, a model must *bootstrap* itself to discover viable solutions. Our SDS curriculum is more than just a technical achievement in that it is a step towards aligning artificial and biological motor control.

Indeed, the SDS policy exhibits a number of properties that have been observed in primates. First, we found that the SDS policy, without any explicit constraint, discovers a low di-mensional posture and control space, reminiscent of humans performing the Baoding ball task ^3^. While the specific synergies are not the same, it is worth noting that the SDS policy has only been trained on this single task. Second, we found that the controller is robust to activity perturbations and that low-variance PCs still contain task-relevant signals, akin to Yan et al’s experimental result ^29^. Third, we found lower tangling of the dynamics in the learned controller than in the action space, akin to what Russo et al. ^30^ found for the motor cortex vs. EMG dynamics. We observed this last result in a different task and architecture, suggesting it might be a general characteristic of RL policies ^53^.

Our policy analysis further illuminates the complexities of motor control. Low-dimensional control emerges both in SDS and human subjects. Yet, the dimensionality of control in muscle-space and joint-space offers a nuanced perspective, suggesting that assessing complexity solely from motion observations might be misleading ^5,46–49^. This highlights the importance of delving deeper into the underlying representations of control, beyond observations.

### Are muscle synergies an emergent property of successful control?

The terms motor synergies, muscle synergies and motor primitives have been used with diverse connotations in the neuroscientific literature ^5,45,48^. The concept of motor and muscle synergies gained popularity when studies on the frog’s spinal cord showed that the forces generated by stimulating individual sites of the spinal cord combine according to vector summation when the sites are stimulated together ^54^. Analogous results were found in spinalized rats ^55^ and through the electrical microstimulation of the motor cortex of macaques ^56^. These findings suggest that motor commands might result from a linear combination of muscle activation patterns, forming a *basis* (in the algebraic sense) of the motor control space, defined at the spinal cord level. These muscle patterns were also referred to as *motor primitives* ^57^. Other studies, instead, propose that motor primitives are combined by the human motor control system into complex motion through juxtaposition in time, similarly to letters in a sentence ^58,59^. In this line of work, motor primitives are sometimes called *movemes* (since they are to motion what phonemes are to speech). Whether motor and muscle synergies are a fundamental mechanism of motor control to deal with a large control space, the result of optimizing for a specific task, or explained by other hypotheses is an ongoing debate in the motor control community ^3,5,45,47,48^.

We found that like biological agents ^1–5^, RL trained agents learn to operate in a reduced kinematic (pose) and dynamic (muscles) space. However, our dimensionality estimates are larger than in a classic behavioral grasping study by Santello et al. ^43^, and closer to estimates for corresponding object-manipulation tasks by Todorov ^44^. Taking advantage of the biomechanical simulator, like others for different purposes ^5,46,48^, we showed evidence that for successful control one needs many more dimensions than suggested by classic signal reconstruction methods (20 vs. 5). Our analysis is consistent with prior analyses of isometric tasks ^47,49,60^.

One popular explanation for this dimensionality reduction is that it provides evidence that the nervous system might simplify the control problem ^3,5^. We remark that the dimensionality reduction for our RL policies was not imposed by the design of the policy network, but was instead discovered purely through structured learning via the maximization of the reward, reminiscent of a hypothesis proposed by Loeb ^5^. The poor transfer of motor synergies from one task to another (Figure S5) reinforces this interpretation. In fact, the same neural network architecture, trained with the same RL algorithm to control the same hand model, but on different tasks, learns to use different muscle synergies. Our results with reinforcement learning are also consistent with arguments based on optimal feedback control ^10,44,60^. Overall, our results speak against the necessity for muscle synergies as a simplifying constraint to tame the complexity of the biomechanical system. In other words, motor synergies are a signature of good control reflecting plant (biomechanical) and task properties, while not providing evidence for a general simplifying strategy.

### Is curriculum learning necessary for RL?

Despite recent successes in machine learning, optimizability remains a key concern for deep learning, and many innovations seek to target this challenge, such as LSTMs ^61^, ResNet’s skipped-connections ^62^, reward shaping ^63^ and sensorimotor priors ^53,64^, the true potential of RL lies in its adaptability. Our SDS curriculum, inspired by human learning paradigms, curriculum learning ^33,34^ and elements of deliberate practice ^36,38^, exemplifies this adaptability, offering a structured learning pathway that aligns with biological motor control. The success of curriculum learning, mirrored by other top-performing models for the 2022 MyoChallenge, underscores its efficacy ^28^. We note that, A.S.C and A.M. as well as Alessandro Marin Vargas won the MyoChallenge in 2023^65^, with a solution also relying on curriculum learning, suggesting that it is a necessary ingredient for mastering complex skills with reinforcement learning, despite recent advances in exploration and reward shaping for musculoskeletal systems ^42,66^. Thus, we currently believe curriculum learning is necessary for acquiring complex musculoskeletal skills. Additionally, self-play ^67,68^, a framework in which an agent competes against itself, can be regarded as a form of implicit curriculum.

### Do humans use curriculum learning for skill acquisition?

Practice is essential for learning motor skills, and the order in which one practices greatly impacts success ^6,37,38,69^. Importantly, humans can learn (certain) skills in a few trials without requiring a curriculum ^37^. We speculate that for Baoding balls in particular, humans already have extensive prior experience about manipulating objects with their hand and thus will rely on a mechanism akin to transfer learning. This differs from the learning framework we used to train the SDS policy network, which starts learning from scratch (i.e., randomly initialized weights). The artificial model that starts from random weights can be considered as a purely exploratory agent without prior knowledge. This scenario aligns more with the acquisition of complex or “unnatural” skills (like learning to do a backflip), or at early development stages, situations in which humans greatly benefit from structured practice depending on one’s skill level ^35–38^. SDS provides evidence for the utility of those ideas in a challenging reinforcement learning setting. At the brain level, practicing with increasing task difficulty based on the individual’s skill level gives rise to greater performance and increased corticospinal excitability ^70^, compared to practice at constant difficulty. Going forward, there are many exciting questions at the interface of skill learning and motor neuroscience. Advances in pose estimation ^71^, biomechanics as well as curriculum learning, as presented here open up new possibilities. For instance, how do EMG patterns change when humans learn a novel, complex motor skill?

### Limitations

Our study has several limitations. First, like any biomechanical model, the MyoHand is an imperfect, simplified imitation of a human hand ^72^. As all the environments in MyoSuite, it does not consider some features of the human musculoskeletal system, such as the elasticity of the tendons and the pennation angles of the muscular fibers (see Methods). Furthermore, anatomical components such as the skin are not part of the model.

Second, the SDS policy network is an abstraction of the sensorimotor system, with important limitations. It provides a general sensorimotor transformation, that receives sensory feedback as an input, processes it with recurrent transformations and then projects it to muscles activations. Its design is similar to what was used in previous modeling studies ^30,50,51^, but it is of course not a reflection of the known anatomy of the motor system. For instance, it is not hierarchical and does not include reflexes ^5,9,13,73^.

Third, the type of sensory feedback provided to the SDS policy is different from the one received by the motor control system. In fact, in the Baoding balls task (as part of the NeurIPS competition), proprioceptive feedback is provided in joint coordinates, which is unbiological ^39,40^. As the SDS policy was part of a competition, we were not allowed to change the input signals, which had to be the same for all the participants. We provided evidence that SDS can also be trained with feedback of muscle lengths, muscle velocities and muscle forces (Figure S2). Future work should consider providing the control policy with proprioceptive information more closely resembling the output of muscle spindles and Golgi tendon organs ^5^. Last but not least, the Baoding balls environment directly provides the state of the balls to the policy network, in the form of their instantaneous position and velocity. In this way, touch and vision are bypassed. While inferring the state of the balls from those senses is in principle doable, it is beyond the scope of this study. Furthermore, besides achieving a higher score than the other solutions, the SDS policy required 40% less energy than the closest competitor ^28^. However, future work could explore how patterns of muscle recruitment and metabolism relate to humans.

In conclusion, our work demonstrates the potential of realistic biomechanical simulation-based approaches in motor control research. We showcased several techniques that help us understand the workings of the artificial motor control agent, such as control subspace inactivation and inter-tasks tangling metrics across the processing pipeline. Of course, this analysis departed from the existence of a successful artificial motor policy in the complex Baoding Task, which we achieved by combining a coaching-inspired curriculum with Deep RL. The result of the training was a successful motor policy that, while it significantly differs from its biological analog, retains important aspects of it. Crucially, as the gap between the artificial and biological policies closes, analysis techniques like the ones in this paper will offer increasingly powerful insights to complement experimental findings in biological motor control. As we continue to push the boundaries of musculoskeletal simulations in RL, the quest for model alignment in sensorimotor systems remains at the forefront, promising a future where the intricate movements of living organisms can be seamlessly replicated and understood in the digital realm.

## Acknowledgments

We thank members of the Mathis Group for helpful feedback. A.M. is appreciative to the Kavli Institute for Theoretical Physics (KITP) in Santa Barbara, where part of the manuscript was written. AM thanks Nicola Hodges and John Krakauer for discussions on skill learning.

## Funding

A.C. and A.M. are funded by Swiss SNF grant (310030_212516). A.I. acknowledges EPFL’s Summer in the Lab fellowship to join the Mathis Group. A.M. was supported in part by grants NSF PHY-1748958 and PHY-2309135 to the KITP. P.T. and N.P. were supported by University of Geneva internal funding.

## Author contributions

A.S.C., P.T., N.P., A.P. and A.M. conceived SDS. A.S.C., P.T., N.P. wrote the code for winning the competition and compared performance to baselines. A.S.C and A.M. conceived the analyses. A.S.C carried out all analyses with initial help from A.I. A.M., A.S.C., P.T., and N.P. wrote the manuscript with input from all authors. A.M. and A.P. supervised the project and acquired funding.

## Declaration of interests

The authors declare no competing interests.

## Methods

Code for training RL policies (https://github.com/amathislab/myochallenge) and model weights as well as rollouts are provided (https://zenodo.org/records/13332869). Code used for analyses in this paper is available in the following repository: https://github.com/amathislab/MyoChallengeAnalysis.

### Musculoskeletal model of the forearm (MyoHand)

The simulation environment of the Baoding balls task is part of the library MyoSuite ^26^. The musculoskeletal model of the forearm is called *MyoHand*, previously used in the library MyoSim ^72^. MyoSim is a library of biomechanical models ported to the MuJoCo physics simulator ^27^ from models in OpenSim ^14^. MyoSuite defines environments based on these models (that also include reward functions, etc.), where policies can be trained with RL. MyoHand comprises of 29 bones, 23 joints and 39 muscle-tendon units. It is based on widely used models of the human hand (*2nd-Hand* ^74^) and of the human forearm (MoBL-ARMS ^75^), as implemented in the biologically-accurate simulator OpenSim ^14^. Caggiano et al. ^26^ merged the two models and enhanced them with the addition of an Opponens Pollicis muscle, to obtain a complete model of the human forearm apt for object manipulation. Differently from OpenSim, MyoSuite is implemented in the MuJoCo physics simulator, which enables faster execution by up to three orders of magnitude. Compared to Open-Sim, MyoSuite adopts a simplified muscle model, which, e.g., does not consider tendon elasticity and fiber pennation angles.

The muscle activation dynamics and the force-velocity relationship used in MyoSuite are identical to those used in OpenSim. The muscle models of MyoSuite obtain accurate force-length-velocity curves, by optimizing the available parameters in order to match the more detailed model of Open-Sim ^72^. This optimization process happens in three steps: 1) Matching forward kinematics, which ensures that the joint and limb positions align accurately with biomechanical data from OpenSim 2) Matching moment arms of each muscle, which verifies that the leverage effects of muscles across joints are consistent with OpenSim, and 3) Matching force-length validation curves, which adjusts the muscle models to replicate force generation behaviors as observed in Open-Sim. Wang et al. ^72^ report a relative root mean squared error (RMSE) for the muscle moments of 0.38 ± 0.57% and of 4.1 ± 2.0% compared to the OpenSim model. This Open-Sim model was chosen as the benchmark for the MyoHand as it provides a highly accurate reproduction of the biological moment arms for all the intrinsic and extrinsic muscles of the hand (1.5 mm average RMS error across the moment arms of all muscles, and 7.1% relative error between artificial and biological muscle attachment points ^74^). Importantly, however, OpenSim still cannot model all the details of a human hand, and further improvements in our understanding of motor control may come from improvements in the accuracy of the musculoskeletal models.

### Baoding balls challenge

The interaction between the control policy and the MuJoCo physics simulator can be formulated as a Partially-Observable Markov Decision Process (POMDP) ℳ= ⟨𝒮, *𝒜, 𝒪, 𝒯*,, *γ, ⟩*. The process ℳ is identified by the state space 𝒮, the observation function 𝒪, the action space 𝒜, the transition function 𝒯, the reward function ℛ and the discount factor *γ*. The action space 𝒜 ⊂ ℝ^39^ is the space of the possible activations of the 39 muscles controlling the human arm model. The observation function 𝒪: 𝒮 → ℝ^86^ maps the state to the observation vector, which includes the angular position of each joint (23 elements), the positions (6 elements) and velocities (6 elements) of the two balls, the positions (6 elements) and the distances (6 elements) of the targets and the activation of each muscle at the previous time step (39 elements). The transition function 𝒯: 𝒮 × 𝒜→𝒮 maps a state and an action to a new environment state, defining how the environment evolves depending on the agent’s decision. An agent seeks to maximize the discounted cumulative reward 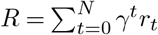, where *r*_*t*_ is the reward at time step *t*, associated by the reward function ℛ: 𝒮 ×𝒜 ×𝒮 → ℝ to the transition from the state *s*_*t*_ to the state *s*_*t*+1_ via the action *a*_*t*_.

The performance of the agent for the Baoding ball challenge (Figure 1G,H) is evaluated in terms of the *solved fraction* (*SF*), corresponding to the number of steps in which both balls are in proximity of the two moving target balls:

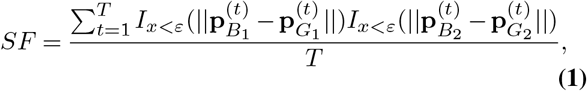

where *T* is the maximum number of time steps in the episode, *I*_*·*_(*x*) is the indicator function, *ε* is the distance threshold between the balls and the targets (1.5 cm), while 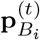 and 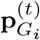,*i* ∈ {1, 2} correspond to the position vector of the ball and the goal *i* at time step *t*, respectively.

Phase I of the MyoChallenge had the objective of rotating the two balls counter-clockwise, with a constant rotation period of 5 seconds. The simulation was chunked into episodes of 200 time steps, corresponding to 5 seconds of simulation. This means that the agent had sufficient time to complete a full rotation of the two balls, before the simulation terminated. An early termination condition was set to accelerate the training, when the balls fell below the palm. In this way, no computation was wasted simulating the environment with the balls in a hopeless position (namely, too far from the hand), which would not provide any useful experience. At the beginning of every episode, the balls started from the same initial location, corresponding to the first target position.

Phase II of the MyoChallenge introduced many complications compared to Phase I. At the beginning of every episode, a random task was sampled (clockwise or counter-clockwise rotation, holding the balls in place) and a target trajectory generated accordingly. Differently from Phase I, the starting point of the target trajectory might be different from the ini-tial position of the balls. This meant that in Phase II it was impossible for the model to score 100% *solved*, because in most episodes the targets would spawn away from the initial position of the balls, and the model would require some time steps before being able to reach them. This complications were addressed with several intermediate curriculum steps (Figure 1I, Phase shift). Besides the random task selection, further variability was provided by the randomization of the task parameters, namely: target rotation period, ellipse axes of the rotation trajectory, mass, size and texture of the balls. Each of these parameters was sampled independently, uniformly at random in a predefined range (Table 1). The MyoChallenge is described in Caggiano et al. ^28^.

### Reward engineering

The performance score (Eq. 1) provided a too sparse signal to be directly optimized via reinforcement learning, as an agent which had not yet learnt how to rotate the balls would almost always collect a vanishing score. For this reason, we designed a dense reward function ℛ: 𝒮 → ℝ, which associates a meaningful performance score to all states. The reward was computed as a weighted sum of four values: the distance between each ball and the corresponding target (2 values), the indicator function representing whether both balls are at most 1.5 cm away from the respective target and the indicator function representing whether the balls are still on the palm. This last reward component proved fundamental for the agent to learn not to drop the balls, which would cause an early termination of the episode (and thus a lower cumulative reward). We penalized the distance between the balls and the targets (weight: -1) and promoted the balls being on the palm (weight: 1) and close to the targets (weight: 5).

### Reinforcement learning details

We used the on-policy RL algorithm PPO ^18^ from the Stable Baselines 3 library ^76^ with a recurrent architecture that has LSTM layers ^61^ in both the actor and critic, which allowed us to deal with the partially observable environment (Figure S1). The neural networks of the actor and the critic were implemented in PyTorch ^77^. The parameters of PPO and of the network architecture are listed in Table 2. This policy was trained as a baseline. To win the challenge we developed the following curriculum learning strategy.

### Static to Dynamic Stabilization curriculum

The schematic of the SDS training curriculum is illustrated in Figure 1F. For clarity we only show the CCW trials, but the full curriculum includes CW and Hold trials. The key idea of the SDS curriculum is to learn stable postures at intermediate states along the desired trajectory *before* having to learn how to reach those states. The benefits of learning stable intermediate postures are twofold: (1) they serve as safe arriving and departing subgoals for an increasingly complex and unstable policy and (2) they robustly shape the value function of the RL agent such that it assigns a high value to these intermediate states, effectively acting as attractors for the final policy. The SDS curriculum imposes several subtasks that the agent must solve before moving to the next subtask. In the first subtask, the balls are initialized at random phases along the desired rotation cycle, and the goal of the agent is simply to hold them still at the initial position (Figure 1F, first panel). In the following subtasks, the balls are also initialized randomly, but now the task of the agent is to move them following the desired trajectory, gradually increasing the target speed from subtask to subtask. As the curriculum advances and the targets speed up, at one point it is not beneficial to use random initialization anymore, as the policy can benefit from exploiting the inertia of the balls. At this point, SDS initializes the balls at the original initial position of the task (Figure 1F, second to last panel) and continues speeding up the targets until it reaches the final task.

The code for training the policy including all hyperparameters is available at online (see link above).

### Control task

To contextualize the analysis of the Baoding policy, we considered the policy trained by Chiappa et al. ^42^ for the Hand Pose task of MyoSuite ^26^. The environment uses the same musculoskeletal model of the forearm as the Baoding balls environment, but rewards the agent when it reaches a certain target pose (i.e., an angular position of each joint). At the beginning of each episode, a new (randomly sampled) target pose is selected. Each target joint angle is sampled independently from the others, uniformly at random in half of the attainable range of the corresponding joint, in a neighborhood including the resting position. This makes the task hard, as some pose configurations might not be achievable due to mechanical coupling. As for Baoding, the training also relies on PPO and recurrent networks, with the same network architecture (Figure S1).

One episode includes 100 simulation steps, corresponding to 2.5 s of simulation time. Each episode starts with the hand in a resting position and with a new target pose. The reward function is maximized when the hand achieves the target pose as fast as possible and maintains that pose for the longest time. The function is an equally-weighted sum of three components:

- *pose* corresponds to the euclidean distance between the hand pose and the target hand pose.
- *solved* assigns a positive reward when the root mean squared error (RMSE) between the hand pose and the target hand pose is smaller than 7.34 degrees.
- *penalty* assigns a negative reward when the RMSE between the hand pose and the target pose is larger than 57.65 degrees.

The used policy achieves a solved fraction of 68.3%, meaning that the hand successfully maintains the target pose for more than 2/3 of the simulation time.

### Additional tasks: Hand Reach, Reorient, Pen

To evaluate the similarities between control policies trained on different tasks, we considered the Hand Reach, Reorient and Pen environments from MyoSuite, and in particular the policies trained by Chiappa et al. ^42^. All these environments feature the same musculoskeletal model (MyoHand). These policies present the same architecutre as all the other policies analyzed in this paper, and were trained with the same RL algorithm (PPO). We refer to ^42^ for the details of the training setup, while we provide a description of the tasks below.

Hand Reach is a free movement task, which does not involve object manipulation. The objective of the policy is to control the MyoHand and reach five target points, one per finger tip, at the same time and as fast as possible. The target positions are sampled at the beginning of every episode within a range specified by the environment. One episode includes 100 sim-ulation steps, corresponding to 2.5 s of simulation time. The policy achieves 0.654 *solved fraction*, meaning that it manages to keep the finger tips at the target positions 65.4% of the time. Reorient and Pen, on the other hand, are object manipulation environments. The policy has to use the MyoHand to move and rotate a die (Reorient) and a pen (Pen) to achieve a target orientation. One episode lasts 150 simulation steps (3.75 s) for Reorient and 100 simulation steps (2.5 s) for Pen. A new target orientation is sampled at the beginning of each episode. The *solved fraction* is 0.685 for Reorient and 0.648 for Pen.

### Dataset generation

For analyzing the SDS policy, we created two datasets including 1000 episodes each, in which the artificial agent trained for the MyoChallenge rotates the two Baoding balls for 5 seconds. In the first dataset, Baoding-SV (small variations), we sampled in each episode the size and the mass of the balls within a small range of values, to introduce some variability in the task while maintaining consistent movements. This dataset is used for the analyses illustrated in Figures 2, 3, 4, 5 and in Figure 6A-B.

The second dataset, Baoding-LV (large variations), features the same experimental conditions as the Phase II of the MyoChallenge. Baoding-LV was used to linearly decode the unobservable environment variables from the policy’s neural population (Figure 6C).

Furthermore, we created a third dataset (Control) comprising 1000 episodes collected with the control policy trained in the Hand Pose environment, where in each episode a new target pose is sampled from the training distribution (although the specific pose was not observed during the training). Data extracted from the Control dataset are utilized in Figure 3A-B.

Finally, we created datasets comprising 1000 episodes using the policy resulting from the twelfth SDS curriculum step and the policies trained on Hand Reach, Reorient and Pen. These datasets have been used to generate the plots in Figures 5 and S5.

In each dataset, we recorded the joint angles, the joint velocities and the muscle activations with a frequency of 40 Hz. Furthermore, we recorded all the internal activations of the policy network, namely, the memory state and the output of the LSTM layer, and the output of the two fully connected layers (Figure S1). We validated the size of the dataset verifying that the dimensionality of the joint angle, joint velocity and muscle activation trajectories was stable when only using a subset of the data (Figure S4).

### Estimation of the number of motor synergies

We analyze synergies as in previous works ^43,44^. The coordination between muscle activations, hand poses or hand velocities was quantified via principal component analysis (PCA). The first principal components capture the (linear) basis that maximally explain the variability of the dataset. We applied PCA on the hand poses, hand velocities and muscle activations of the episodes included in the Baoding-SV, Control and other datasets. For the Baoding-SV dataset, we also analyzed how these quantities change when considering the sub-dataset of episodes where the balls rotate in the same direction (Table 3). For consistency with Todorov and Ghahramani ^3^, we calculated the number of degrees of freedom by averaging the number of principal components necessary to achieve an explained variance higher than 85% and 95%. The same steps were applied to the Control dataset, to estimate the motor synergies emerging from free hand motion (target pose reaching).

### Control subspace inactivation (CSI)

Control subspace inactivation (Figure 4A) is a procedure to evaluate the impact that removing certain muscle synergies has on the task performance. Given a policy *π* : ℝ^*N*^ → ℝ^*M*^, *N* being the size of the observation space and *M* the size of the action space, we consider an orthogonal matrix *W* ∈ ℝ^*M ×M*^, whose columns **w**_*i*_, *i* = 1, …, *M*, define an orthonormal basis of the action space (in this paper, the space of all the possible muscle activations). Given an observation **o** ∈ ℝ ^*N*^, the policy returns a muscle activation pattern **a** = *π*(**o**) ∈ ℝ^*M*^. Control subspace inactivation modifies the muscle activation before it is transmitted to the physics simulator, selecting a subset *I* of the indices of the basis vectors **w**_*i*_, whose contribution to the control signal is removed. The resulting activation pattern is

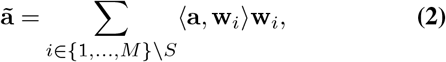

where ⟨ ·, ·⟩ indicates the canonical inner product of ℝ ^*M*^. A reward and a new observation are then returned by the environment, according to the transition dynamics of the POMDP. In the experiments illustrated in Figure 4B, Figure 5C,F and Figure S5, control subspace inactivation used the orthonormal basis defined by the principal components extracted by one of the datasets of this paper, inactivating components in order of importance. When all the components are inactivated (left most point in the plots), the performance is 0, since a constant action is applied at all times. When no component is inactivated (right most point in the plots), the full task performance is recovered.

### Decoding the rotation direction from the PCs of the hand pose

We assessed whether progressively less important principal components retain information about the rotation direction by decoding it via logistic regression. To this end, we considered four different subsets of three principal components of the hand pose, extracted from the Baoding-SV dataset (Fig 4B). One data point of the training dataset consisted in a flattened vector of three principal components of the pose for all the 200 time steps of one episode (600 features in total). The mean accuracy was obtained by averaging the logistic regression score via 5-fold cross validation on random shuffles of the Baoding-SV dataset. We used the Python library Scikit-Learn ^78^.

### Time-dependent importance of the PCs

The relative importance of the PCs of the hand pose and of the muscle activations varies throughout the rotation of the Baoding balls. To visualize the point at which each component is more relevant, we considered the subset of the Baoding-SV dataset featuring clockwise rotations and averaged the hand poses and the actions. In this way we obtained the average hand pose and muscle activation across episodes at each time step. For clearer visualization, these coefficients were then rescaled between −1 and 1 (Figure 2B). A full rotation of the Baoding balls corresponds to two periods in the hand pose and muscle dynamics, which are evident in several principal components. In fact, approximately the same PCs swap the first and the second ball and vice versa, to complete a rotation. The re-utilization of PCs is indicative that the RL policy has achieved appropriate generalization capabilities: if the identity of the balls is ignored in the input (which is irrelevant for task performance), the states at the beginning of the first and second half of the cycle are similar, so the PCs that achieve the desired goal should also be similar.

### Visualization of the network activations

We embedded the observations, the actions and the network activations into a 3-dimensional space using Uniform Manifold Approximation Projection (UMAP) ^79^. Each point of the graphs of Figure 6A corresponds to a single time step.

### Tangling of the population activity at variable depth

We quantified the tangling of the observations, actions and population activity of each layer of the SDS policy (Figure 6A, B) employing a metric introduced by Russo et al. ^30^. They propose to measure the time-dependent tangling of a trajectory **x** : ℝ → ℝ^*d*^ with the scalar function *Q*(*t*), defined as

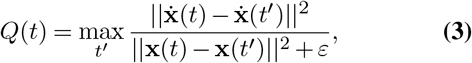

where 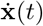 is temporal derivative of the trajectory, while *ε* is a small constant such that division by 0 is avoided (*ε* = 10^*−*10^ in our analysis). The tangling was computed on a lower-dimensional projection of the signal (8 dimensions), obtained via PCA like in Russo et al. ^30^. A single value per plot (Q value in Figure 6A) is computed by averaging *Q*(*t*) over *M* trajectories and *N* = 200 time steps:

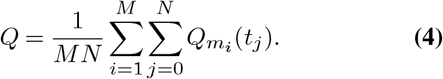

We also report Q values per trial (episode) by averaging across *N* = 200 time steps (Figure 6B).

### Decoding the environment variables from the population activity

Some unobservable (i.e., not part of the observation provided to the agent) parameters of the environment, such as the mass of the balls, have an impact on the dynamics of the environment and thus on the task performance. Such parameters are encoded with variable accuracy in the population activity of the different layers of the SDS policy. Using the Baoding-LV dataset, we computed the encoding score of each environment parameter proportionally to how well such parameter could be linearly decoded from each layer of the policy network (Figure 6C). For continuous parameters (Mass, Size, Friction, Radius, Joint Positions, Joint Velocities) we considered the coefficient of determination *R*^2^ of a linear regression, while for the only categorical parameter (Task) we considered the mean accuracy of a logistic regression. To compute robust scores and estimate their variability, we used 5-fold cross validation on random shuffles of the Baoding-LV dataset, using the Python library Scikit-Learn ^78^.

## Videos of the policies

Here we provide videos of several episodes in Phase I and Phase II.

**Video 1.**
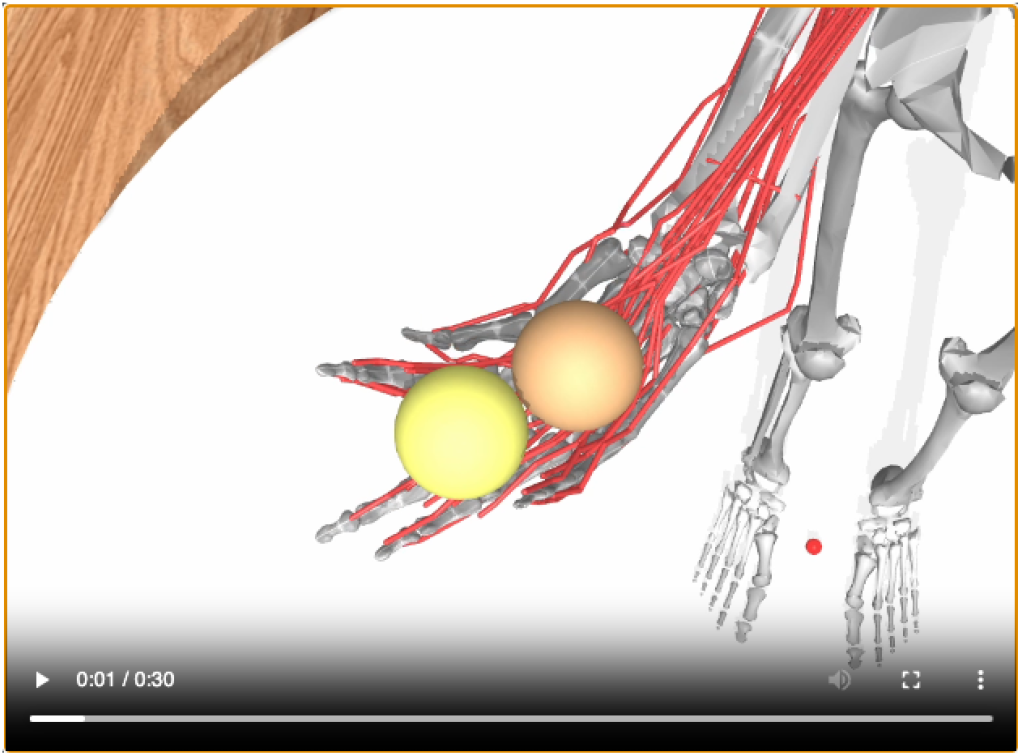
Here: A single frame a video of the policy which scored 100% in PhaseI. The attached video shows a visualization of the hand and ball dynamics. Several episodes were concatenated. This policy was not part of the curriculum, which led to the final policy of Phase II, because it was trained to perform a single rotation direction. For Phase II, we restarted the same SDS curriculum as Phase I, but introducing both rotation directions since the very beginning. Video available at: https://youtu.be/RFIWLEUUqbw

**Video 2.**
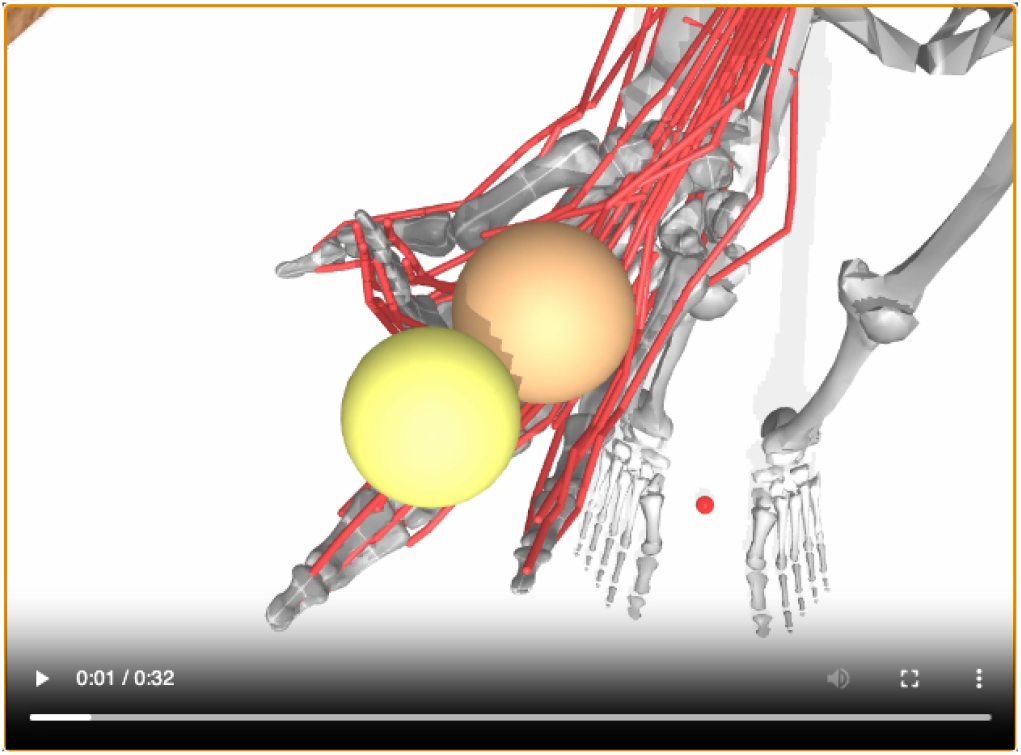
Here: A single frame a video of the policy which scored 55% in Phase II. The attached video shows a visualization of the hand and ball dynamics. Several episodes with different hyperparameters and conditions (CCW, CW, hold) were concatenated. The sometimes visible small balls indicate the target position. When the Baoding balls are both close to the target positions, they turn brighter. The score is given by the fraction of time steps in which such condition is satisfied. Video available at: https://youtu.be/fF72wf0JK-4

## Supplementary Figures

Here we provide a diagram of the policy and critic network (Figure S1), the performance of SDS when trained based on more realistic proprioceptive feedback (Figure S2), dynamics per joints and muscles for one episode (Figure S3), a validation of the size of the dataset (Figure S4) as well as our analysis on the generalization of muscle synergies from multiple tasks to the Baoding Ball task (Figure S5).

**Figure S1.**
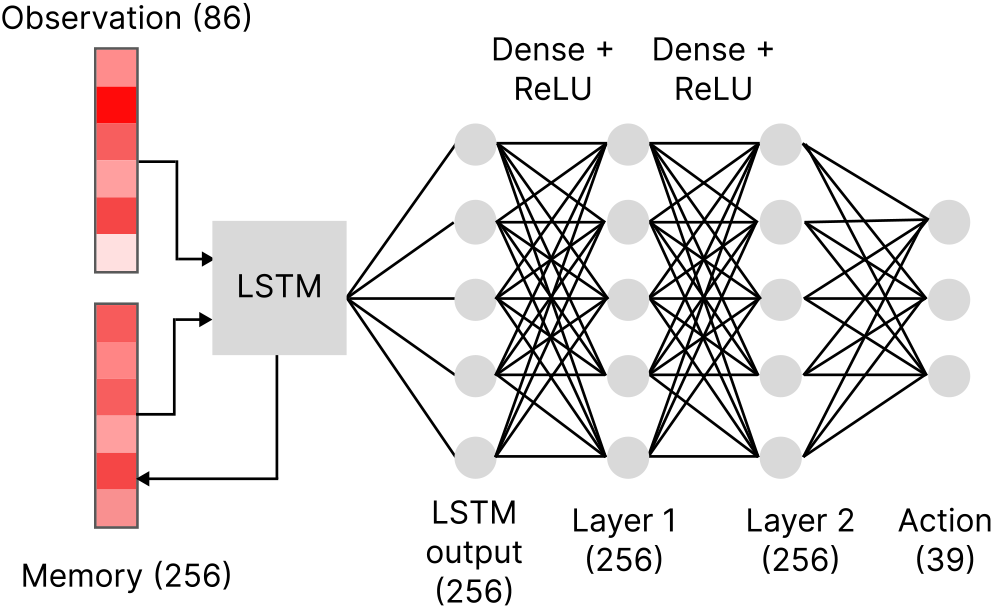
Architecture of the policy network, related to Figures 1 and 4 and Methods. The observations are fed into a single LSTM layer, followed by two fully connected layers from where the action is predicted. Both the actor and the critic have the same architecture. The numbers between parentheses indicate the size of each layer.

**Figure S2.**
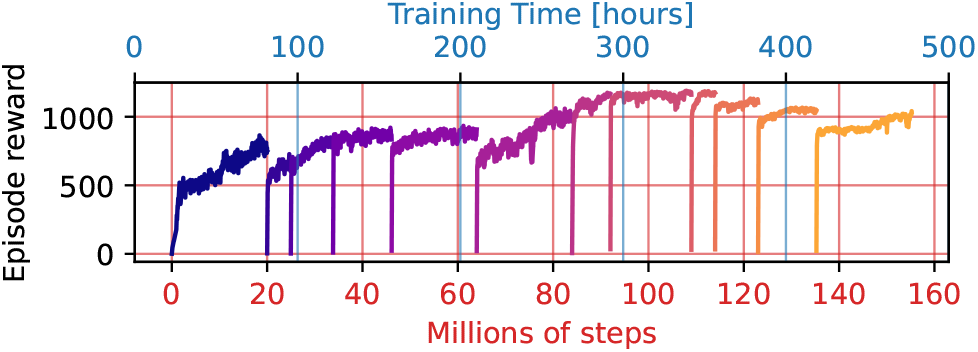
SDS curriculum with muscle observations, related to Figure 1. Learning curves of Baoding Phase I, replacing the hand pose in the observation with muscle lengths, muscle velocities and muscle forces. Also with the modified observation format the SDS curriculum successfully solves Phase I, achieving over 99% solved. In the interest of time and as this is a proof of principle, we did not train this model for Phase II.

**Table S1.**
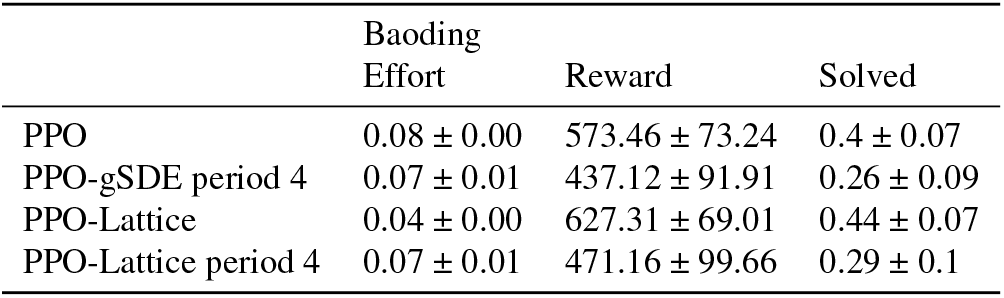
Performance of policies trained on Baoding Phase I with different exploration noises, related to Figure 1. Effort is a measure of the total muscle activation, while Reward and Solved are performance metrics (the higher the better). Results are adapted from Chiappa et al. ^42^ with Generalized State-Dependent Exploration (gSDE) described in Raffin et al. ^41^. The results are reported as mean ± s.e.m. across 5 random seeds.

**Figure S3.**
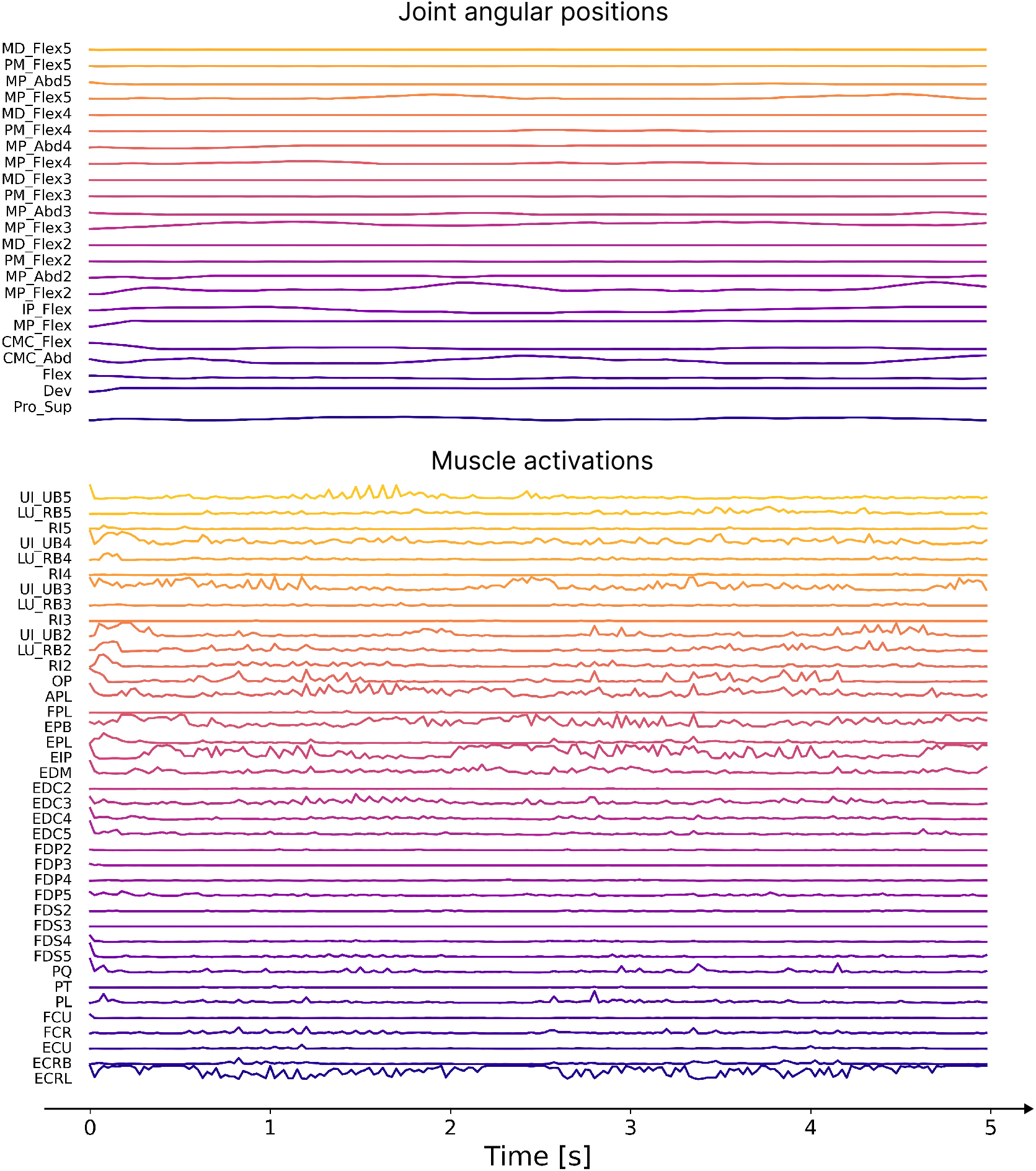
Kinematic and dynamics motifs per joint and muscle, for one full rotation of each Baoding ball, related to Figure 2. The labels of the joints and the muscles are defined according to this convetion: Pro = pronation, Sup = supination, Dev = radial-ulnar deviation, Flex = flexion, Abd = abduction, CMC = carpometacarpal joint, MP = metacarpophalangeal joint, IP = interphalangeal joint, PM = proximal phalanx, MD = middle phalanx, ECRL = Extensor Carpis Radialis Longus, ECRB = Extensor Carpis Radialis Brevis, ECU = Extensor Carpi Ulnaris, FCR = Flexor Carpi Radialis, FCU = Flexor Carpi Ulnaris, PL = Palmaris longus, PT = Pronator teres, PQ = Pronator, EIP = Extensor Indicis Proprius, EPL = Extensor Pollicis Longus, EPB = Extensor Pollicis Brevis. Related to Figure 2.

**Figure S4.**
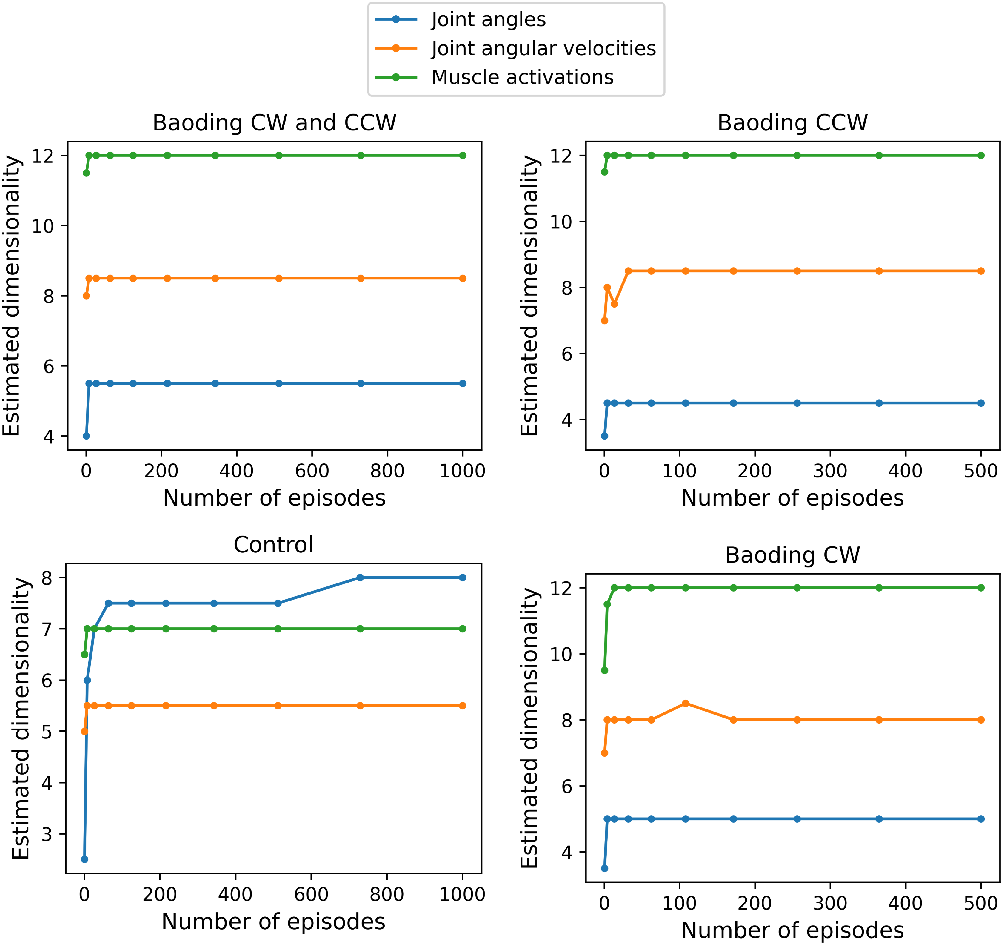
Dataset size validation, related to Figure 3. Number of independent degrees of freedom (as in Figure 3B) vs. dataset size (number of episodes). For all the quantities whose dimensionality we estimated in the analysis, the number of episodes in the Baoding-SV dataset is sufficiently large to achieve stable values.

**Figure S5.**
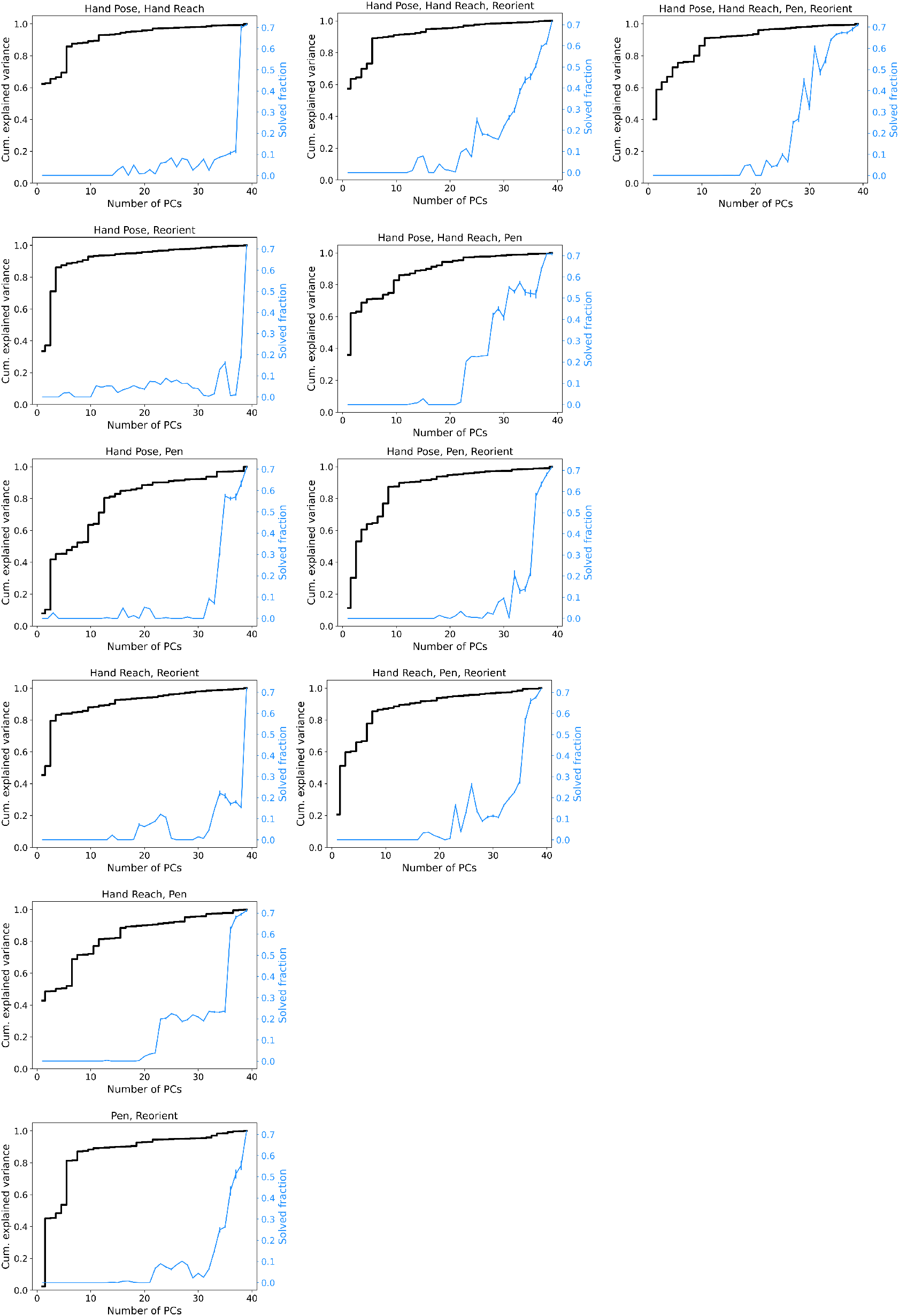
Transfer of muscle synergies from multiple tasks to the SDS policy, related to Figure 5. Cumulative explained variance (black) and task performance (blue) as a function of the number of principal components. The two curves were obtained through control subspace inactivation. Each graph considers the principal components extracted from policies trained on a different combination of tasks, with the tasks listed in the title. From left to right more tasks are combined. Surprisingly, the performance only increases beyond 20 dimensions, even though a substantial fraction of the variance is already captured.

## Notes

https://github.com/amathislab/myochallenge

https://github.com/amathislab/MyoChallengeAnalysis

https://zenodo.org/records/13332869

### Competing Interest Statement

The authors have declared no competing interest.

### Summary of Updates

Substantially updated article; muscle synergy analysis for multiple tasks.

